# A Biophysical Model for ATAC-seq Data Analysis

**DOI:** 10.1101/2024.01.25.577262

**Authors:** Catherine Felce, Gennady Gorin, Lior Pachter

## Abstract

The Assay for Transposase-Accessible Chromatin using sequencing (ATAC-seq) can be used to identify open chromatin regions, providing complementary information to RNA-seq which measures gene expression by sequencing. Single-cell “multiome” methods offer the possibility of measuring both modalities simultaneously in cells, raising the question of how to analyze them jointly, and also the extent to which the information they provide is better than unregistered data where single-cell ATAC-seq and single-cell RNA-seq are performed on the same sample, but on different cells. We propose and motivate a biophysical model for chromatin dynamics and subsequent transcription that can be used with multiome data, and use it to assess the benefits of multiome data over unregistered single-cell RNA-seq and single-cell ATAC-seq. We also show that our model provides a biophysically grounded approach to integration of open chromatin data with other modalities.

## 1 INTRODUCTION

The Assay for Transposase-Accessible Chromatin using sequencing (ATAC-seq), introduced in 2013, has become a widely adopted epigenetic discovery tool that can be used to identify regions of open chromatin (1). In 2015, this method was extended to allow for measurements at single-cell resolution (scATAC-seq) (2, 3). These techniques were initially developed and applied independently from transcriptomics experiments, and ATAC-seq data are still often analyzed in this manner, albeit often by re-purposing gene-expression tools (4). As a result, several methods have emerged for analyzing ATAC-seq data in isolation (5, 6).

However, since the major interest in ATAC-seq data lies in elucidating the gene expression programs within cells, the data are best understood in combination with gene expression data. The integration of bulk ATAC-seq and RNA-seq data has been carried out in various forms, e.g. (7) and (8) (spatial). However, approaches thus far have been based on heuristics and do not consider underlying biophysical processes such as transcription factor binding, transcription bursting, processing and splicing of transcripts, and degradation. ATAC-seq and RNA-seq data can, in principle, inform the specifics of models for these processes. These insights can then be fed back into mechanistic models, which can be used to analyze data and provide information about the dynamics of regulation and transcription.

Approaches for integrating unregistered scATAC-seq and scRNA-seq data, i.e. data collected from different cells from the same sample, include joint analysis of differentially accessible regions (DARs) (9) and differential gene expression. Such methods have been used to compare gene regulation across tissues between healthy and pathological conditions (10–15), and across tissue types (16), as well as to identify new gene-regulatory pathways (17, 18). Combined use of scRNA-seq and scATAC-seq data can also help to distinguish cell-types (19, 20) and scATAC data can be used to impute scRNA data (21). An exciting recent development has been the ability to perform registered scATAC-seq and scRNA-seq, identifying DNA fragments and mRNA transcripts from the same individual cells (22, 23). Such methods provide a direct avenue for joint analysis of the two modalities, thus avoiding the need for “integration” of scRNA-seq and scATAC-seq (24), albeit raising new challenges including sparsity of registered data (25).

While both registered and unregistered data have been useful for a variety of applications, analysis methods have not incorporated biophysical modeling, and have instead relied on heuristics and phenomenological inferences (20). While progress has been made in building biophysically motivated and interpretable models for RNA-seq data (26–28), and Gorin et al. have outlined a general chemical master equation (CME) formulation for describing various models of transcription with intrinsic and extrinsic noise (29), these tools have not been applied to ATAC-seq, or joint analysis of ATAC-seq and RNA-seq.

Building on (29), we propose a model for chromatin dynamics, inspired by the Ising model from physics. This constitutes a first step towards biologically interpretable joint modeling of ATAC and RNA-seq data (see Figure 1). Using a combination of theory and simulation, we investigate the propagation of gene-gene correlations from the level of chromatin accessibility to the level of transcript counts. Fitting our model to ATAC-seq datasets provides some justification for our characterization of chromatin dynamics. Within our modeling framework, we also consider the utility of registered vs un-registered data, and investigate the value of combined single cell measurements for the development of biophysical models.

**Figure 1.**
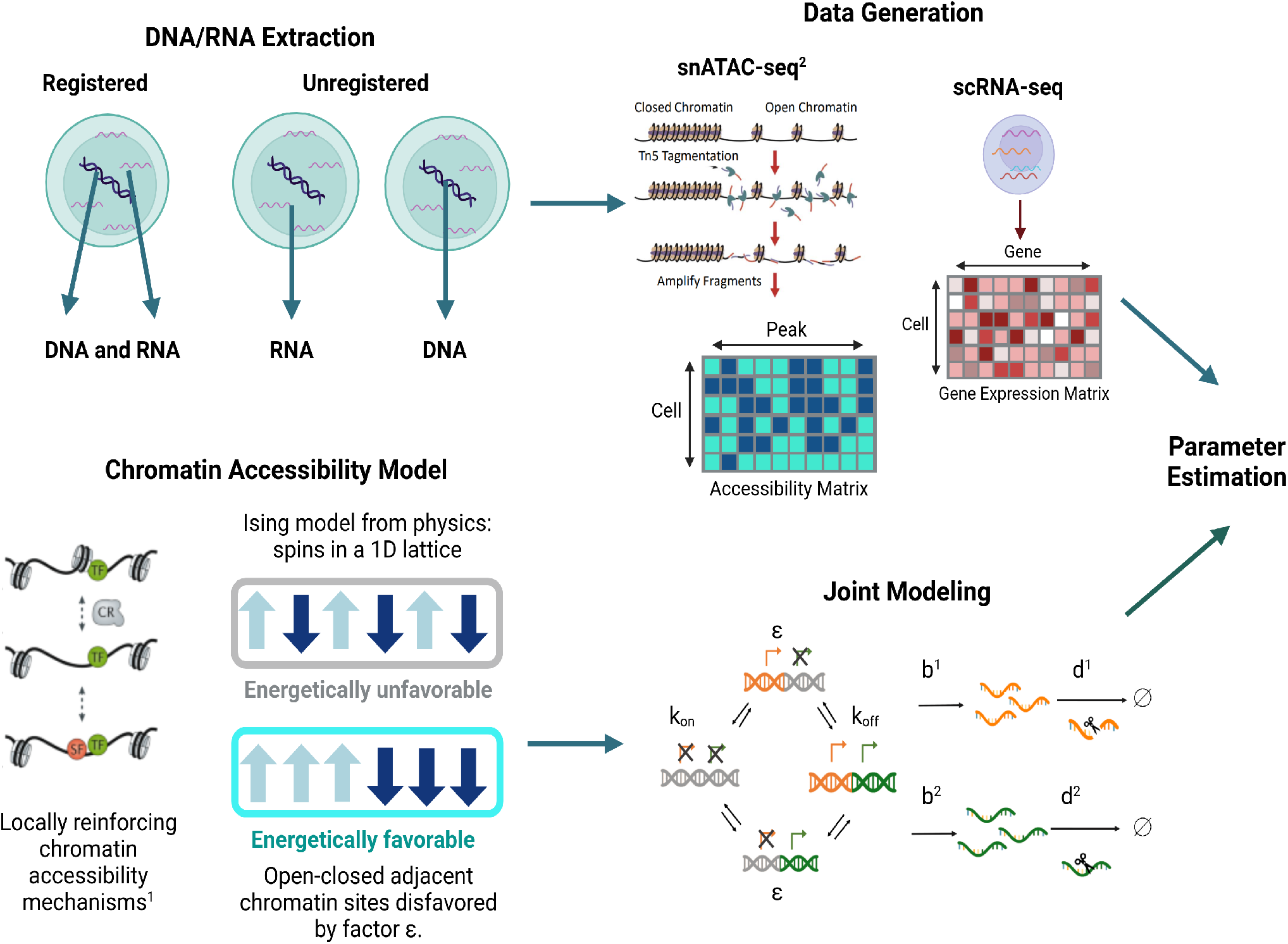
Joint modeling of gene accessibility and constitutive mRNA production allows for parameter estimation using scATAC-seq and scRNA-seq data. **DNA/RNA Extraction:** Registered scATAC-seq and scRNA-seq information can be taken from the same cells in multiome data. Alternatively, scATAC-seq and scRNA-seq can be performed on separate cells within the same sample. **Data Generation:** The transcripts captured during scRNA-seq are sequenced and processed to give a cell-by-gene expression matrix. The fragments of accessible chromatin captured during scATAC-seq are sequenced, and accessible regions are identified via peak calling on the bulk data. Fragments from each cell are then aligned to these peak-regions, giving a binary cell-by-peak matrix, indicating whether or not each cell contained a fragment from each peak region. **Chromatin Accessibility Model:** We choose a model for chromatin accessibility where neighboring peak-regions are correlated within each cell. This is reminiscent of the Ising model from physics. **Joint Modeling:** By combining a Markov chain model for chromatin-state switching, (illustrated, left), with assumptions for subsequent RNA dynamics, (e.g., constitutive transcription at accessible regions), we can estimate the joint distribution of transcript counts and chromatin accessibility across cells. The model is parameterized in terms of RNA characteristics for each transcript species (e.g., transcription rate: *b*^*i*^, decay rate: *d*^*i*^, for the *i*^*th*^ species), and features of the chromatin-state transition matrix (e.g. neighbor-neighbor correlations: *ϵ*, rate of gene switching on/off: *k*_*on*_/*k*_*off*_). **Parameter Estimation:** By comparing the observed data with predicted distributions from the joint model after adding noise, we can estimate parameters. References ; 1: (30), 2: (31)

## 2 MODEL DETAILS

### 2.1 Chromatin Neighbor Interactions

Recruitment of active chromatin re-modelers by pioneer transcription factors could lead to expanding regions of open chromatin (see stabilizing role of secondary transcription factors in (30)). These mechanisms could bias adjacent DNA regions towards sharing the same state of transcriptional accessibility. This motivates the use of correlation structure to describe chromatin, and we consider the simplest model that could yield such correlation: first order neighbor interactions reminiscent of the Ising model from physics. We choose to parameterize this model by a parameter *ϵ* < 1, which encodes the preference of neighboring chromatin sites to be aligned. *ϵ* = 1 yields site independence, and allowing *ϵ* to differ from 1 gives us the next simplest model, where one misalignment in a configuration reduces the probability of the configuration by a factor *ϵ*. Note that in our parametrization, a smaller value of *ϵ* corresponds to a stronger correlation between neighboring regions. Such a formulation is supported by exploratory analysis, as shown in Section 3.3.

The Ising model from statistical mechanics describes magnetic behavior in terms of individual dipole spins which can each be in either of two opposite directions, denoted as up and down, or +1/*−*1. In ferromagnetic materials, it is energetically favorable for neighboring spins to be aligned. In the presence of an external magnetic field, it is also favorable for spins to align with the direction of the magnetic field, and the interplay of these two factors determines the probability distribution of different states of the system. Such systems can be simulated using Glauber dynamics, where spins are flipped with a probability proportional to (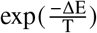), where Δ*E* denotes the change in the energy of the system due to the flip. In our chromatin model, up and down spins are analogous to open and closed regions of chromatin, and our model is therefore effectively the 1-D Ising model.

Ising models have been used to analyze ChIP-chip data and discover transcription factor binding sites (32, 33). Mo et al. consider a hidden Markov model (HMM) treating chromosomally consecutive probes in a microarray as neighbors in a 1-dimensional Ising chain. The hidden state of the system is a specific configuration of enriched vs non-enriched probes in the chain. Although their treatment and the connection to the Ising model is similar to ours in this work, the foundation for their Ising hypothesis is technical rather than biological, as it rests on the correspondence of many consecutive enriched probes to a single protein binding event.

In the Ising model, the external magnetic field is assumed to be constant for all spins in the lattice. However, inspection of the first order moments for chromatin accessibility from ATAC-seq data suggests that this feature of the model is not appropriate in our context (see Supplementary Note S1.2, Figures S1, S2, S3). Therefore, we allow the ratio of chromatin opening and closing rates to vary between sites, giving a separate ‘field strength’ parameter per site, plus one correlation parameter. This gives us, for example, a seven-parameter model to describe the chromatin aspect of the biological system for a six-site locus (before considering technical noise).

### 2.2 CME Implementation

To implement our model we follow the Chemical Master Equation (CME) approach of Gorin et al.(29).

The system is assumed to be Markovian, and the probability distribution across states evolves in a deterministic manner. The state of the system is described by the number of transcripts present for each gene, and the state of chromatin openness for each gene. In our analysis, we have assumed a one-to-one mapping between ATAC-seq peak-regions and genes. We assume constitutive transcription whenever a gene is in its open (or on-) state, and no transcription in the closed (or off-) state.

The processes of gene-switching, transcription and decay occur at given rates, and these rates are the parameters of the system. These processes each contribute terms to the system of ordinary differential equations (ODEs) which govern the time evolution of the system. For example, decay processes contribute terms to the ODEs which move probability from states with higher transcript numbers to states with lower transcript numbers. These decay terms are straight-forward and affect states independently from their chromatin configuration. In contrast, transcription rates at each gene can depend on the underlying state of the DNA.

To illustrate the CME framework, we give the simple example of a single-gene system: the telegraph model introduced by Peccoud and Ycard (34). The chromatin configuration of the system consists of the single gene being either open or closed for transcription, with corresponding transcription rates of *b* or 0 respectively. The transcript decay rate for this species is *d*, and the rates of chromatin opening/closing are *k*_on_ and *k*_off_ respectively. The system would then be governed by the following ODEs:

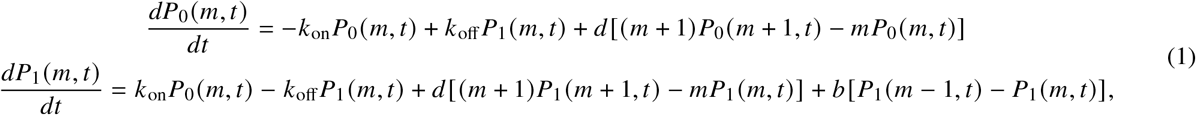

where *P*_*α*_ (*m*, ***t***) represents the probability of the system to be in the gene-state indexed by *α*, where here *α* = 0, 1 corresponds to the gene being closed/open for transcription respectively, with an integer number, m, of transcripts, at time t. Note that, despite the use of a full derivative, the value of *m* is fixed in each equation, and Equation 1 represents a system of ODEs, one for each possible value of *m*. The first terms encode switching between gene-states, whilst the decay and transcription terms represent changes in transcript number. The transcription terms only affect the evolution of probabilities in state 1, since no transcription occurs in state 0.

These equations can be summarized in a matrix form. We introduce a system probability vector ***P***(*m*, ***t***), whose components are the *P*_*α*_ (*m*, ***t***) described above. The gene-state dynamics can then be encapsulated in the transition matrix, H, given by:

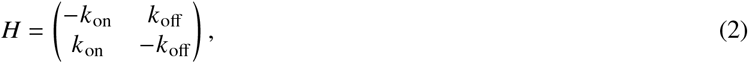

and the evolution of the system can be succinctly described via:

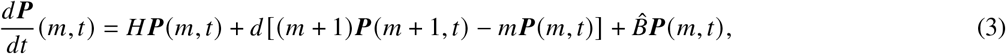

where we have also introduced a diagonal transcription matrix, 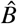, defined such that 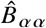 is the transcription rate in the state indexed by *α*. In this case, 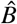 is given by:

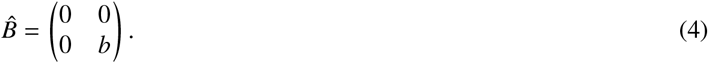

This construction can be extended to multiple transcript species and a greater number of gene states. Although we consider only those chromatin-state transitions which can be effected through the opening or closing of a single gene, the framework allows for an arbitrary graph structure of transitions, via modification of the transition matrix.

In the following analysis, we consider a space of 64, (2^6^), gene-states corresponding to a binary openness value for six different genes, and vectors ***m*** of length six, where each vector index corresponds to the integer number of transcripts for a different gene species.

#### 2.2.1 Chromatin Dynamics

Working within the CME framework of (29), we consider the gene state to evolve in a Markovian manner according to the evolution equation:

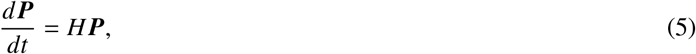

where ***P*** is a vector denoting the probability to be in each chromatin configuration. We can then encode cooperation between neighboring regions within the gene-state transition matrix, H.

We define a chromatin-state matrix *S*, such that 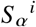 gives the openness of gene *i* in chromatin-state *α*, and 1/0 correspond to open/closed genes respectively. Note that here and in what follows, for matrices we use superscript Latin indices to reference DNA sites (genes), and subscript Greek indices to reference chromatin configurations. In this work, we will consider the chromatin-state indexed by *α* as a string of 1’s and 0’s, representing the openness/closedness of adjacent DNA sites. This gives 2^*n*^ configurations for *n* DNA sites. As a toy example, we consider a system of only two genes, which would have the state matrix:

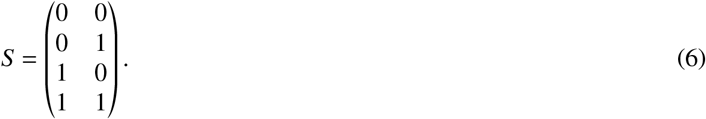

For this two-gene system we can choose the gene-state transition matrix:

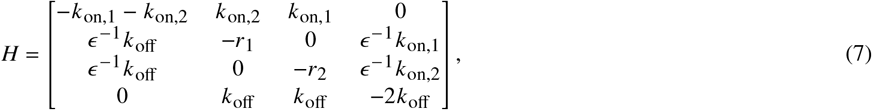

where *r*_*i*_ *≡ ϵ* ^*−*1^(*k*_on,i_ + *k*_off_), and *∈* is defined as in Section 2.1.

Note that to reproduce the Ising stationary distribution for the case of varying site-openness propensities, we need to set *k*_off_ as equal for all sites, and allow *k*_on_ to vary between sites. This gives a stationary distribution where the probability of each state is proportional to the product over sites of each *k*_on_ value, and *∈* to the power of the number of ‘misaligned’ neighbors in the state. (See Section 2.5.)

Working within the CME framework allows straightforward extension of this work to the joint analysis of chromatin accessibility and transcriptomics.

#### 2.2.2 RNA Transcript Count Evolution

We define the probability generating functions (PGF) with gene state index *α* as:

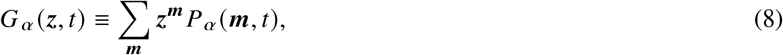

where *P*_*α*_ (***m***, *t*) represents the probability of the system to be in the state indexed by *α* and have *m*^*i*^ RNA counts of each transcript *i*, and ***z*** represents the vector of PGF arguments, [*z*^1^, …, *z*^*n*^], for RNA species 1 to *n*. We define 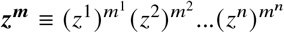.

Let ***G*** represent the vector of PGFs, [*G*_0_, …, *G* _*N −*1_], for gene states 0 to *N −* 1. Let *B* be the production matrix, whose entries 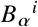 represent the rate of production of species *i* in state *α*. For the simplest model, where transcript *i* is produced at rate *b*^*i*^ for states where gene *i* is active and at rate zero otherwise, we have the relation: *B* = *S*diag(***b***), for ***b*** the vector of transcription rates with components *b*^*i*^. Let ***d*** be a vector representing the decay rates of species *i*. Following the procedure outlined in Gorin et al.(29), we then have the full evolution equation:

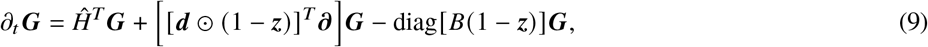

where the Hadamard product between two matrices is (*X ⊙ Y*)_*αβ*_ *≡ X*_*αβ*_*Y*_*αβ*_ for any two matrices *X* and *Y* with the same dimensions (here dimension *n*×1 for *n* the number of RNA species). (1 *−* ***z***) is the vector with components 1 *− z*^1^, 1 *− z*^2^, …, 1 *− z*^*n*^, and the second term on the right-hand side of Equation 9 involves a sum over partial derivatives with respect to PGF variables *z*^*i*^.

### 2.3 Moments and Statistical Properties

The PGF evolution equation above leads to the following moments and statistical properties, before considering noise.

#### 2.3.1 Gene-State Moments

For considering moments, we introduce the vector ***π***, representing the steady-state probabilities of the chromatin states of the system. The vector ***π*** has the length of the entire state space, i.e. 2*n*, where *n* is the number of sites in the locus. To obtain the steady-state, we set the left-hand side of Equation 5 to zero. In terms of the chromatin-state transition matrix, *H*, this gives:

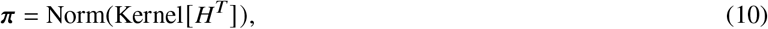

where ***π*** is normalized such that its entries sum to one. We denote the chromatin-state vector by *σ*, where *σ*^*i*^ represents the openness of the *i*^*th*^ chromatin region. (For the gene-state indexed by *α, σ* is equivalent to row *α* of the state matrix, *S*). We consider a first moment of the system, the average openness value of an individual DNA site, *i*. From the state matrix *S*, defined above, we have:

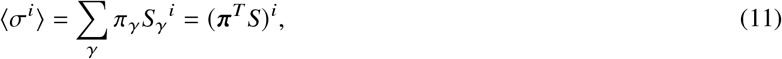

where the ⟨⟩ denote expected values and *σ*^*i*^ represents the openness of the ith region. The gene-state covariance in this notation is given by:

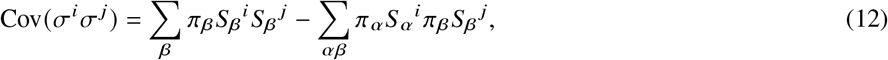

where the variance can be obtained by setting *i* = *j* in this expression.

#### 2.3.2 RNA Transcript Count Moments

To find moments concerning the *i*^*th*^ RNA transcript species, we differentiate Equation 9 with respect to *z*^*i*^ (see Supplementary Info Section S4.2 for details). This gives the transcript means for each species *i*:

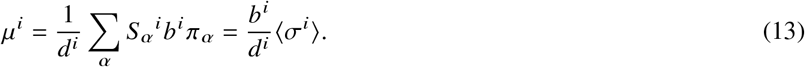

Defining the matrix:

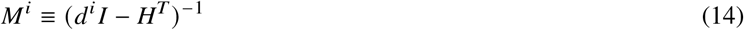

we arrive at an expression for the transcript covariances:

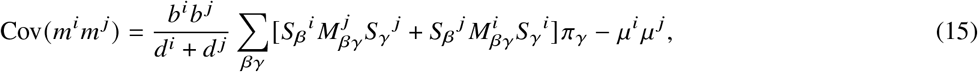

and variances:

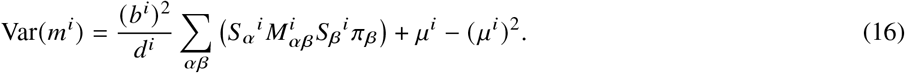

#### 2.3.3 Correlation Propagation

We define site-site correlations for accessibility and transcript count respectively via:

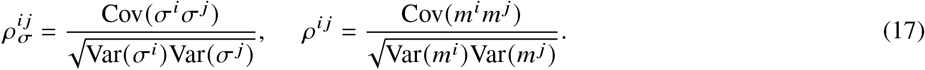

We define *f* as the ratio between these correlations:

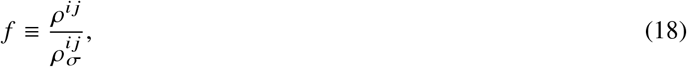

and an expression for *f* is given in Supplementary Info Section S4.3, along with its value for a range of parameters in Figure S5. We might assume that correlations between the transcript numbers for different genes would be weaker than the chromatin-level correlations between the parent genes, given that transcript dynamics are downstream of chromatin-state dynamics. We might also expect that the correlations at the transcript and chromatin levels would have the same sign. These constraints together would suggest that *f* should be contained within the range 0 < *f* < 1. However, *f* can exceed 1 for certain parameter ranges in our model, (as visible in Figure S5). To explore the intuition behind this result, see Supplementary Note Section S4.4. We define two illustrative toy systems, with | *f* | > 1 and *f* < 0 respectively, (see Figure S6), and confirm the behavior of their correlations in Figure S7.

### 2.4 ATAC-seq Noise Treatment

For data analysis, we added noise to the result of the above CME dynamics in the following way. We start from the steady-state gene-state distribution ***π*** defined in Equation 10. Then, with the addition of technical noise, we reach a final steady state distribution, 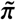, arrived at via:

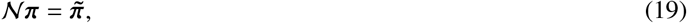

where we have defined a noise matrix, 𝒩. We consider the properties of this noise matrix for dropout noise which symmetrically affects all sites in a locus, giving them a probability *p*_drop_ of being lost. Although this technical noise model is, in a sense, symmetrical, there is an asymmetry in the space of binary strings, since transition from a state with fewer on-sites to a state with more on-sites is impossible. In this sense, the possible states of a locus form a partially ordered set. For instance,

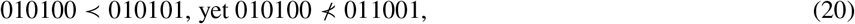

where the partial order, *a ≺ b*, indicates that sequence *a* can be obtained, starting from *b*, via technical noise. In this example, the first two strings are connected by a transition encoded in 𝒩, but the second two are not: the technical noise process cannot create false positives.

In the case of a three-site locus the matrix N is 8-dimensional. Using the basis defined in the Supplementary Note Equation S11, we have:

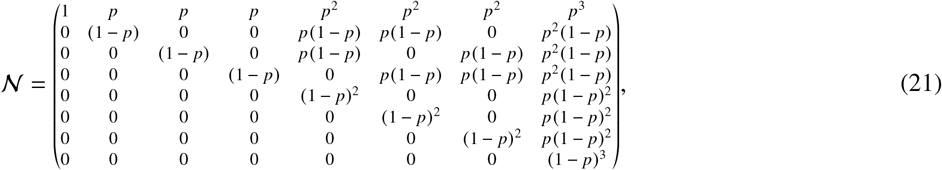

where, here, p = p_drop_, and we have assumed that dropout at each site is independent. This is a triangular matrix. See Figure S4 for a visualization of the effect of this type of noise.

### 2.5 Model Inference

To assess the Ising model, we compare a model fixing *ϵ* to be one, and a model allowing *ϵ* to vary as a parameter. We compare the models using the Bayesian Information Criterion (BIC), which quantifies goodness-of-fit whilst penalizing extra parameters (see Equation S47).

Note that, from above, the probabilities for a chromatin configuration ***σ*** in this model, before technical dropout, are given by:

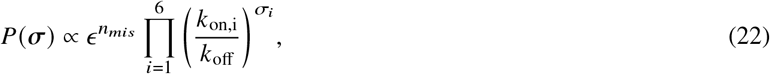

where *n*_*mis*_ is the number of pairs of misaligned neighbors, and again *σ*_*i*_ = 0, 1 for site *i* closed/open. E.g.

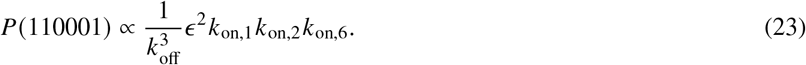

We then add technical noise in the form of binomial dropout, where each site independently has probability *p*_drop_ of flipping from *σ*_*i*_ = 1 to *σ*_*i*_ = 0. However, in the *ϵ* = 1 case, when fitting based on ATAC-seq data alone, this is equivalent to a change in *k*_on_ values. This is because we can express the probability of a site being accessible in terms of its biological *k*_on,i_ and *k*_off_ rates, and multiply this value by (1 *− p*_drop_) to account for technical dropout. The resulting probability of measuring accessibility at a site *i*, given dropout, is then equivalent to the probability without dropout, given the substitution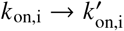, where:

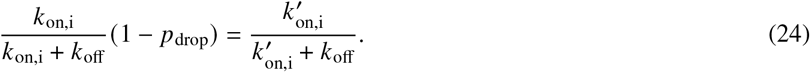

Therefore, after constraining *ϵ* = 1, the reduced model can be fully described using only six parameters (*k*_on,i_ for *i* = [1, 6]). So we are left with two models. In the simple six-parameter model, the loss of reads is absorbed into the ‘field-strengths’, *k*_on,i_ at each of the six sites, i, and the probabilities are given by setting *ϵ* = 1 in Equation 22. This model has independent sites, and thus ignores gene-gene correlations. The model including site-site correlations is given by allowing *ϵ* to vary in Equation 22 to give the initial probabilities for each configuration, and then adjusting these by adding binomial loss at each site. This model does capture correlations between genes, and is reminiscent of the Ising model from physics. The comparison between these two models as applied to ATAC-seq data is shown in Section 3.4.

## 3 APPLICATION TO ATAC-SEQ DATA

### 3.1 Datasets

We use three 10x Genomics datasets to perform our analysis:

- 10k Human PBMCs, ATAC v2, Chromium Controller,
- 8k Adult Mouse Cortex Cells, ATAC v2, Chromium Controller,
- 10k 1:1 Mixture of Human GM12878 and Mouse EL4 Cells, ATAC v2, Chromium Controller.

### 3.2 Data Processing

We pre-processed the single-cell ATAC-seq data using the snATAK pipeline (25). For multiple lanes in a single dataset, we called peaks individually on each lane, merged the peaks and aligned reads to the merged set of peaks. More details of the pre-processing of individual datasets are provided in the Supplementary Info Section S5.

After filtering, we binarized the cell-by-peak matrix. This means that if a cell had at least one read aligning to a certain peak, that peak region was considered open for that cell.

Since our model considers regions which are adjacent in space, we then restricted our focus to those peaks which were within 1.5kbp of their adjacent peak. This distance was chosen somewhat arbitrarily to admit a reasonable number of allowed groups of peaks. In particular, we examined groups of six consecutive ATAC peaks, where each was at most 1.5kbp from the next. Here and in what follows, we refer to these groups of six contiguous peak-regions as ‘loci’.

The distribution at each locus was then calculated by counting the frequency of different configurations across cells. The possible configurations are given by the length-six binary strings, e.g. 110000, which would represent two consecutive open regions followed by four consecutive closed regions.

### 3.3 Exploratory Analysis for Ising-like Model

After selecting suitable genetic loci for analysis, we investigated whether positive correlations are observed between neighboring chromatin sites. We calculated the Pearson correlation coefficient, *r*_*xy*_, between sites in each pair of adjacent ATAC-seq peak regions in the selected loci. (See Supplemental Information Section S1.1 for the details of this calculation). In figure 2, we plot these correlation coefficients against the average openness of the sites, *μ*_av_, for each pair of adjacent sites, across the three datasets. *μ*_av_ is calculated by taking the average openness of a pair of adjacent sites, and then averaging this value over all cells. This exploratory analysis lends support to the use of an Ising-like model, since a consistently positive correlation coefficient between adjacent sites is indeed observed. The proportions of pairs with positive Pearson correlation are 137/150, 35/35 and 68/70 for the PBMC, adult mouse cortex and human-mouse mixture datasets respectively.

**Figure 2.**
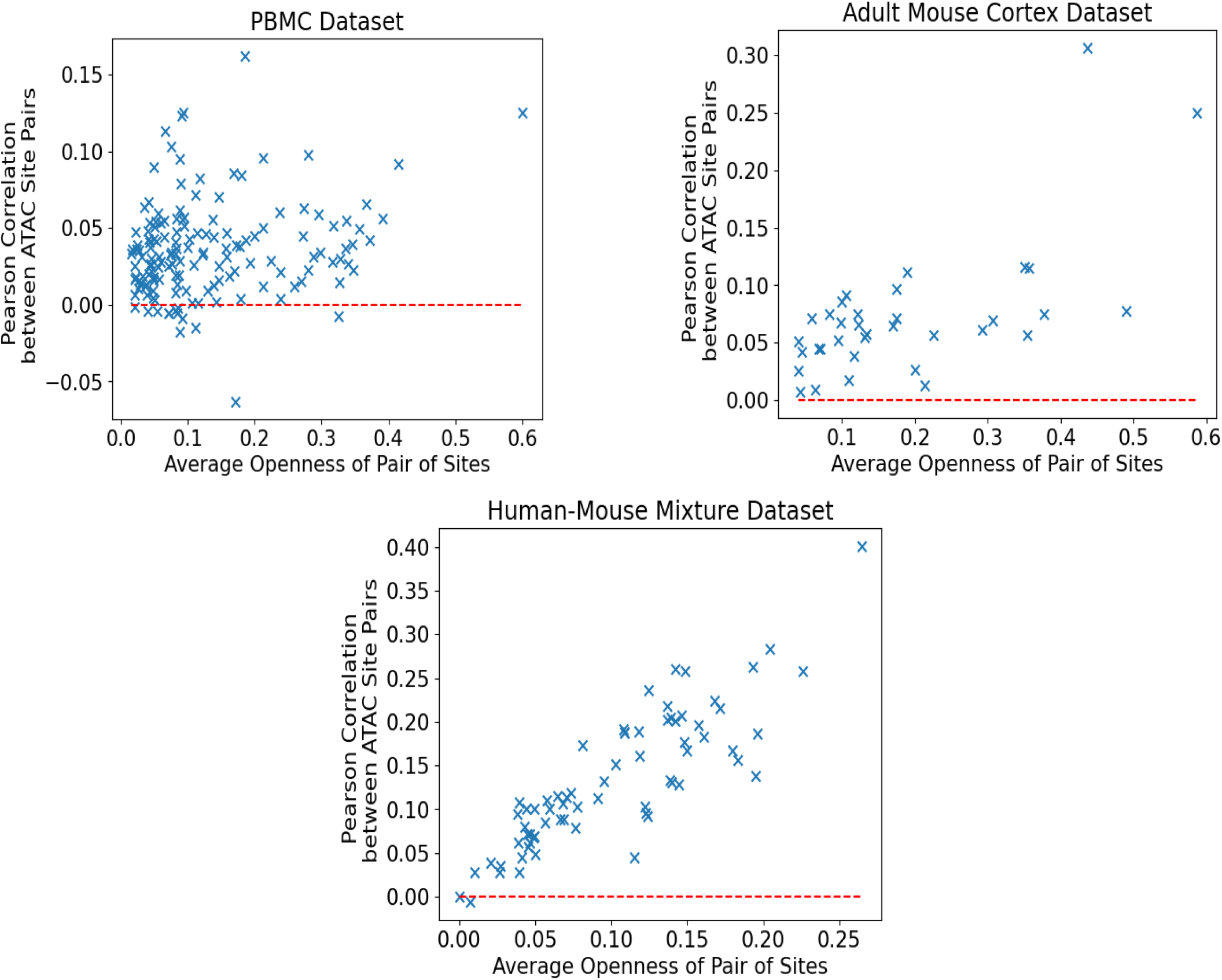
Pearson correlation coefficients between sites in each pair against the average openness of the sites in that pair. Each point represents a pair of adjacent ATAC-seq peak regions included in our selected loci. The red dotted lines are positioned at a Pearson correlation of zero, so all points above the red line have a positive Pearson correlation, lending support to an Ising-like model. **Top Left:** PBMC dataset: 137 of 150 pairs have positive Pearson correlation. **Top Right:** Adult Mouse Cortex dataset: all 35 pairs have positive Pearson correlation. **Bottom:** Human-Mouse Mixture dataset: 68 of 70 pairs have positive Pearson correlation.

### 3.4 Results

After identifying the relevant loci for each dataset, we fit both the six-parameter and eight-parameter models at each locus (see Section 2.5 for a description of the models). We used stochastic global optimization to minimize log-likelihood, and repeated each fit ten times to check convergence.

We show example fits for the six and eight-parameter models in Figure 3, for one six-site locus. The locus is from chromosome 18 of the 10k Human PBMCs, ATAC v2, Chromium Controller dataset.

**Figure 3.**
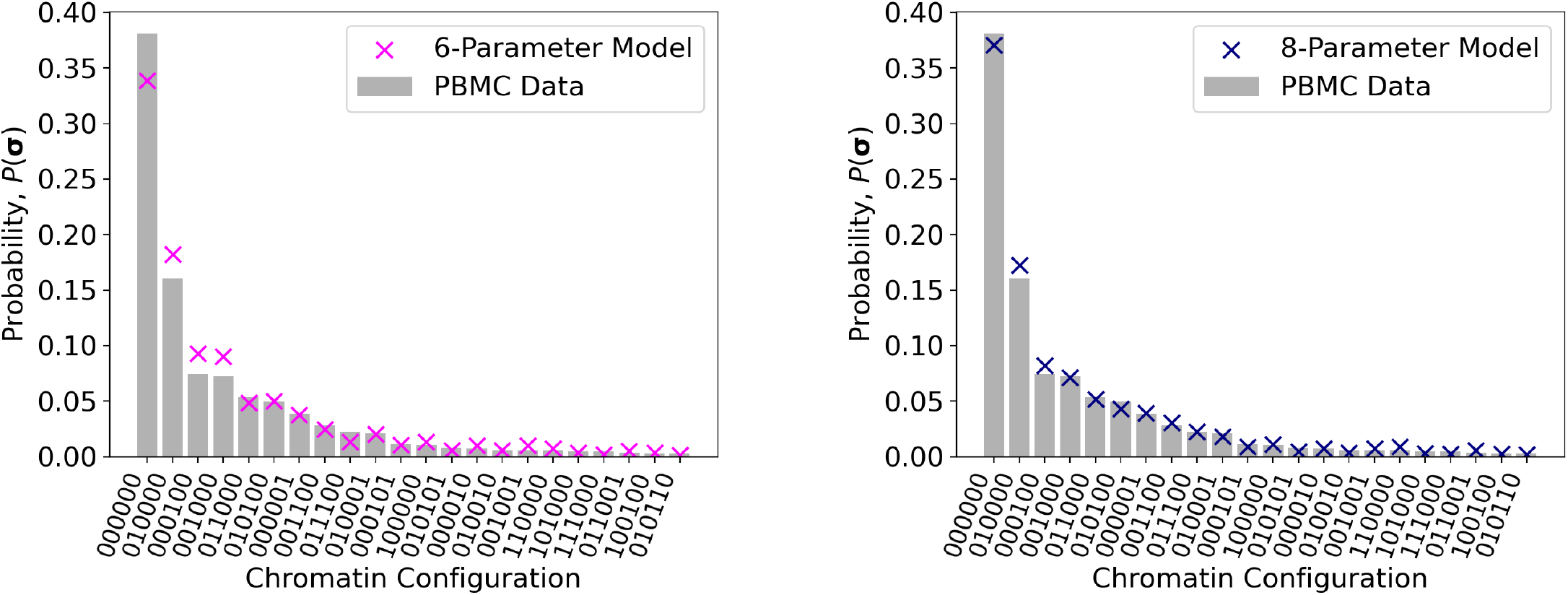
6-parameter (**left**) and 8-parameter (**right**) fits to observed distribution at locus chrom 18:77059667-77072738 from human PBMCs. N.B. The 22 most frequently observed strings of the total 64 are shown on the x-axis. The gray bars show the empirical distribution, and the magenta and blue markers the fitted distribution for the 6 and 8-parameter models respectively.

The bar chart shows the observed fraction of the most frequently observed chromatin configurations at this locus. The magenta and blue markers show the analytic distribution using the best-fit parameters for the 6/8-parameter models respectively. See the Supplemental Note Figure S8 for fits at other loci. We also verified the reliability of the result of the optimization algorithm with an MCMC fit for one locus (see Supplementary Note Figure S9).

To analyze the effectiveness of the Ising hypothesis, we looked at all of the contiguous six-site loci, as defined above, across three different 10x datasets. We calculated the BIC score for the six and eight-parameter models at each locus, and record the BIC score difference in favor of the extra parameter *ϵ* (combined with *p*_drop_). We show the distribution of BIC score differences for each dataset in Figure 4. Almost all of the loci provide support for using the 8-parameter Ising-like model over a model with independent sites. The proportions of loci with a BIC score favoring the Ising-like model are 26/30, 7/7 and 14/14 for the PBMC, mouse cortex and human-mouse mixture datasets respectively. The particularly striking positive outlier in the PBMC plot is locus 10:69043845-69054726. The high BIC difference is due to the much better fit of the 8-parameter model for this locus, as shown in Section S6.2 of the Supplementary Note.

**Figure 4.**
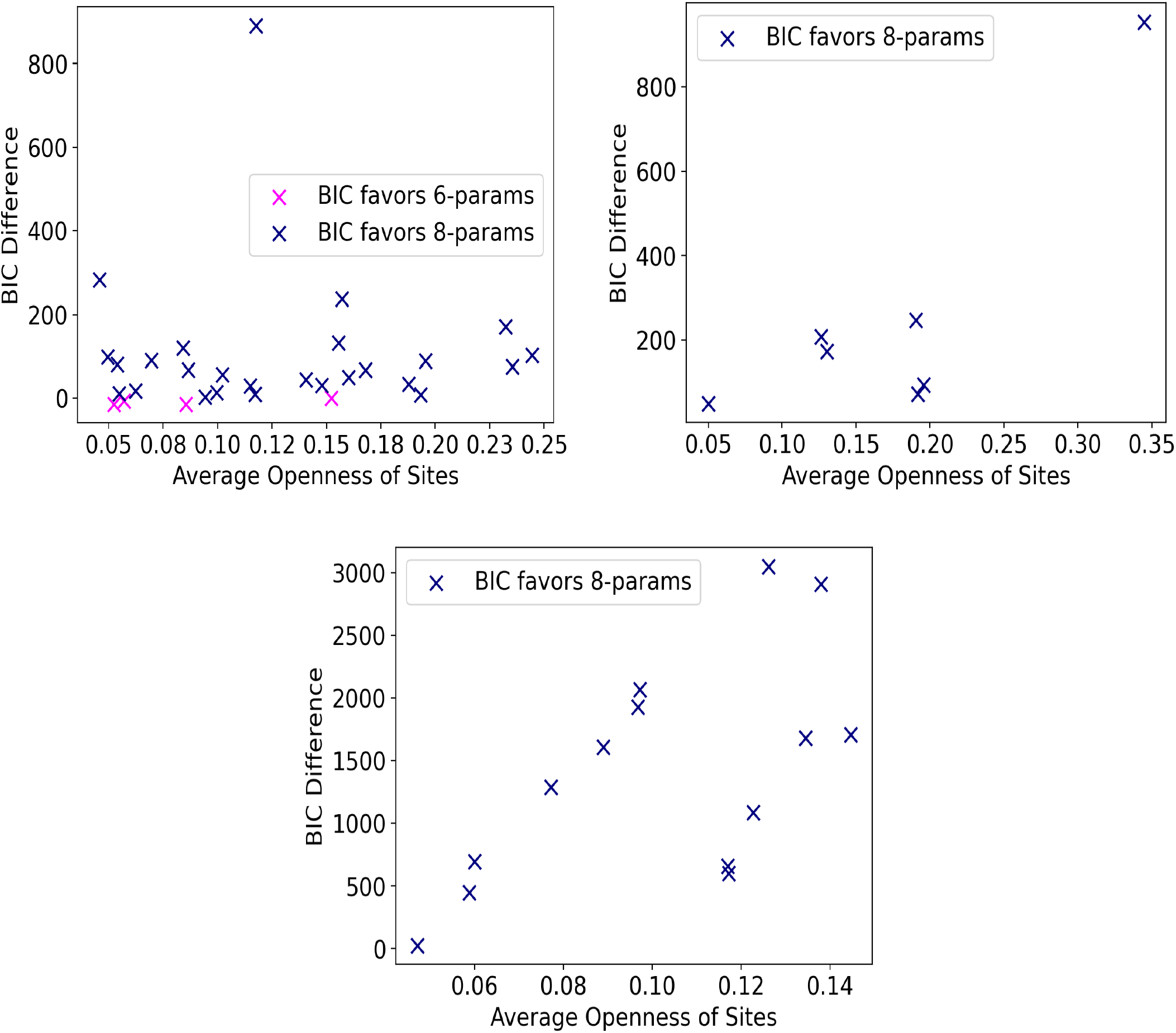
BIC score improvement with inclusion of correlations, i.e. 6-parameter BIC score minus 8-parameter. **Top Left:** PBMC dataset: 26/30 loci support the 8-parameter model. **Top Right:** Adult Mouse Cortex dataset: 7/7 loci support the 8-parameter model. **Bottom:** Human-Mouse Mixture dataset: 14/14 loci support the 8-parameter model.

## 4 DISTINGUISHABILITY COMPARISON: MULTIOME VS UNREGISTERED

For the eight-parameter model described above, we explored the identifiability of the *ϵ* parameter using both registered and unregistered scRNA-seq and scATAC-seq data. In this exercise, we neglect noise.

First we simulated the model described in Section 2.2 for a two-gene system of known *ϵ* and sampled steady-state gene-states and RNA counts for 20, 000 cells. We then solve the system analytically given different values of *ϵ*, whilst keeping the overall probabilities for each gene to be on constant (by adjusting *k*_off_). To calculate the analytic solutions, we numerically solve the PGF evolution equation for the steady state value of the joint probability distribution for gene-state and number of RNA transcripts of each species.

For each analytic solution, we assume that each sampled cell is an i.i.d. variable drawn from this distribution, and therefore that the probability of obtaining this set of simulated data is given by the product of the probabilities for each cell.

For registered data, we calculate the log-likelihood function given by:

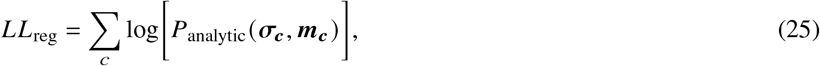

where c indexes cells in the simulated data, *σ*_*c*_ is the gene state observed in cell c (as defined in Section 2.3), and ***m***_*c*_ is the vector of transcript numbers for cell c (as defined in Section 2.2.2).

To show the distinguishability achievable with scATAC-seq or scRNA-seq data alone, we repeat the same process, using the respective marginal probability distributions:

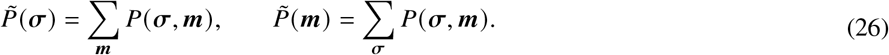

This gives the log-likelihoods:

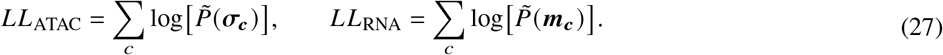

To compare to the unregistered RNA-seq and ATAC-seq data scenario, we sum over log-likelihood contributions from both of the marginal distributions:

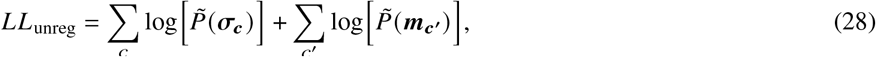

where these two sums are over separate groups of sampled cells.

To assess the information gained from each modality, we make two different comparisons. Firstly, we consider the case where the total number of sampled cells is kept constant. The log-likelihoods in equations eqs. (25), (27) and (28) are calculated for the same 10, 000 sampled cells in each case. For the unregistered case (Equation 28), the first term is a sum over half of these cells, and the second term is a sum over the remaining half. These results are shown for a simulation with *ϵ* = 0.3 in the left panel of Figure 5.

**Figure 5.**
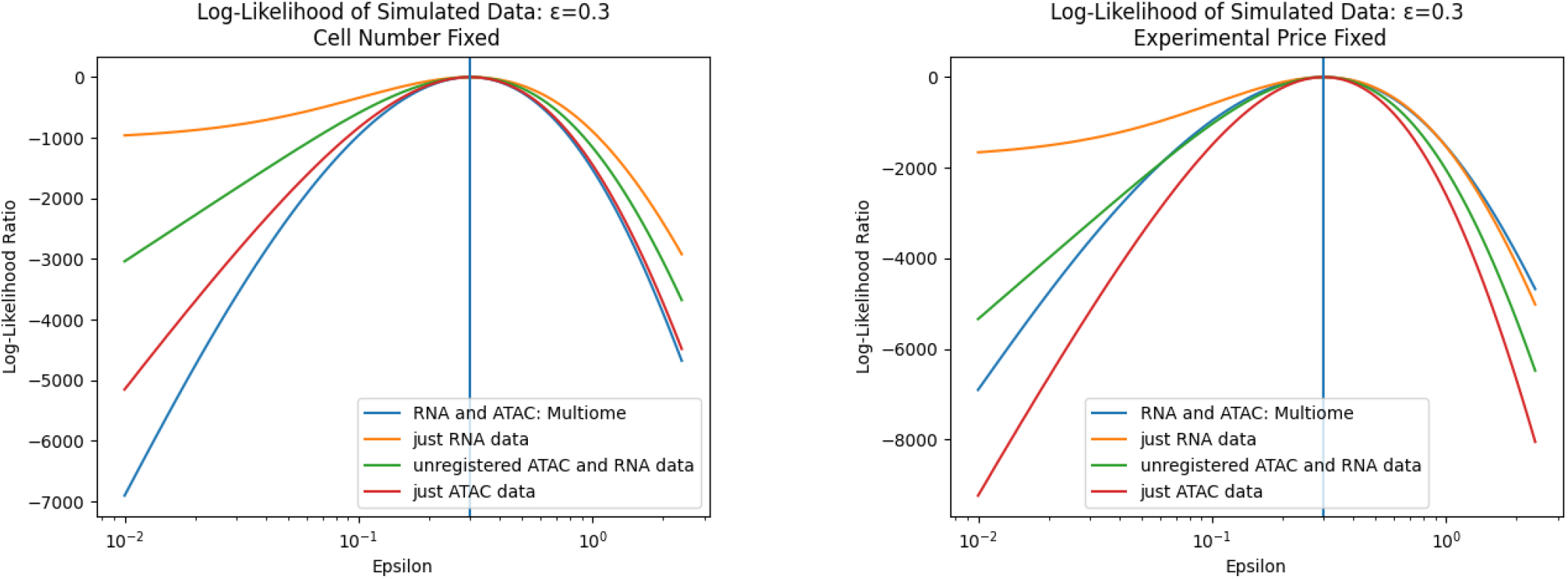
*ϵ* distinguishability for scATAC and scRNA, separately, together and registered, keeping a fixed number of cells (**left**), or a fixed experimental price (**right**).

Secondly, we consider the case of fixed experimental price. We use prices quoted from 10x Genomics for separate scATAC-seq and scRNA-seq vs registered multiome per cell. The ratio of prices per cell was quoted as^1^:

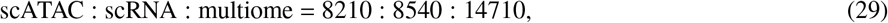

and the price for unregistered scATAC and scRNA is the average of the scATAC and scRNA prices per cell ((8210+8540)/2 = 8375). We then weight the number of cells for each modality by the inverse of its price, using 10, 000 sampled cells for the multiomic case, and proportionally more for the other modalities. The log-likelihood curves obtained via this process are shown in the right panel of Figure 5.

The results in Figure 5 reflect the fact that, whilst multiomic data provides the most distinguishing power per cell for models of this type, (shown by the multiomic log-likelihood curve giving the sharpest peak around the true *ϵ* value in the left panel), after adjusting for price, the results are more complicated. Since the price of obtaining multiomic data is close to the sum of performing scATAC and scRNA separately, unregistered data can sample almost twice as many cells for a similar price. However, information is also lost in the lack of registered RNA and ATAC information from the same cells. The right panel of Figure 5 shows that unregistered and registered multiomic data are comparable in their ability to distinguish *ϵ* in these kinds of models. scATAC-seq data alone would be the most price-efficient method for determining *ϵ*.

## 5 DISCUSSION

In this work we have presented for the first time a minimal biophysical model suitable for joint analysis of RNA-seq and ATAC-seq data. These two modalities are increasingly assayed in single-cell genomics experiments, and our work explores and compares two models which, although relatively simplistic, offer a basic framework for quantitative analysis of multiomic data. We expect that these models will be elaborated on and improved, for example by the inclusion of transcription factor binding. We have already shown that the models help sharpen quantitative questions about gene expression and its relationship to chromatin dynamics, by exploring the ratio of correlations at the chromatin and transcriptome levels.

The framework we have outlined allows us to investigate the effects of DNA-level mechanistic assumptions on the expected distributions of both chromatin configuration and transcriptomics. In this case, we have investigated the effect of neighbor-neighbor correlations on the system dynamics. To undertake our analysis, we have made use of a form of the Ising model from physics, which has been studied at length. Inspired by its use in other areas of computational biology, we have successfully applied the Ising model in the novel context of ATAC-seq data.

After hypothesizing an Ising model for chromatin dynamics, we followed Gorin et al. (29), and encoded these dynamics using chemical master equations. Making simple assumptions about the corresponding RNA dynamics, we were able to formulate the behavior of both modalities in a tractable and biophysically meaningful way. The first and second order moments of the system were simple to derive from the resulting matrix equations, and allowed us to explore the relationship between correlations at the DNA and at the transcript level. In particular, the mathematical tractability of this class of models allowed us to uncover the unintuitive result that downstream correlations between transcripts of different species can theoretically be higher than the correlations of their parent genes at the level of chromatin configuration.

By adding a binomial dropout model for scATAC-seq technical noise, we were then able to directly compare the steady-state distributions derived from the CME formulation to real scATAC-seq datasets. As a proof of concept, we limited our analysis to loci with six contiguous, close ATAC peak sites. We compared a simple six-parameter model of site-independence, with our eight-parameter Ising-like model plus dropout, and found that our model was almost always preferable in terms of the Bayesian Information Criterion. Our analysis highlights the importance of site-site accessibility correlations, despite the sparsity of ATAC-seq data, and informs our understanding of relations between neighboring DNA regions. Our work does not shed light on the biological details underlying these correlations, but the assumptions implicit in our models point towards experiments that may provide more insight into mechanistic underpinnings. For example, loci with BIC scores that highly favor the model including gene correlations could be candidates for more targeted chromatin remodeling or transcription factor binding experiments.

Finally, we used our model to explore experimental design considerations for combining scATAC and scRNA data. In the context of a two-gene system, we have illustrated the ambiguity around the relative merits of multiomic vs unregistered data at fixed price for parameter identifiability. This brings into question the rationale for choosing multiome assays over unregistered data, depending on the question being asked, and also highlights the importance of increased data quality for refining biophysical models. Although this numerical experiment has important limitations – for instance, it omits the impact of dropout noise – it provides a principled foundation for multiomic experiment design. As we obtain a better understanding of technical factors, we can extend this design approach to include noise considerations.

The techniques that we’ve introduced in this paper could be usefully extended to other data-types, such as HiC data. In addition, since our framework couples RNA and chromatin dynamics in a simple and extendable way, our model can be incorporated directly into current biophysical modeling tools such as *biVI* (35). By straightforward modification of our choices for chromatin and RNA dynamics (e.g. constitutive transcription), our approach can be used to explore many more complex systems in a tractable way. Our postulation of an Ising-like structure for the chromatin-state transition matrix could also be extended to analysis of other multiomic assays. Following the prescription outlined in Gorin et al. (29), we could further test the assumptions of this model using single-cell RNA-seq data.

Overall, although sparsity of single-cell ATAC-seq data limits the power of the conclusions reached, we have outlined a promising approach to biophysical modeling of the data type, including in conjunction with single-cell RNA-seq data.

## 6 ACKNOWLEDGMENTS

We thank Charles Trimble for a generous gift to Caltech’s CI2 that partially funded C.F. Thanks to Sina Booeshaghi and Delaney Sullivan for help with snATAK used in pre-processing the single-cell RNA-seq and ATAC-seq data, to Tara Chari and Maria Carilli for helpful feedback on the manuscript, and to Meichen Fang for manuscript feedback in addition to many helpful discussions.

## 7 DATA AVAILABILITY

Scripts implementing these analyses and simulations and reproducing Figures figs. 2 to 5 are available at https://github.com/pachterlab/FGP_2023.

## Footnotes

1 10x Genomics pricing quote

