## Supplemental Note for "A Biophysical Model for ATAC-seq Data Analysis"

### A Biophysical Model for ATAC-seq Data Analysis: Supplementary Information

Felce Gorin Pachter

#### S1 EXPLORATORY DATA ANALYSIS

##### S1.1 Nearest Neighbor Correlations

For each pair of adjacent ATAC-seq peak sites in the loci we selected, we calculated the Pearson correlation coefficient between the two sites in the pair. The sample Pearson correlation coefficient between two variables x and y is defined via:

$$r_{xy} = \frac{\sum_{i=1}^{n}(x_i - \bar{x})(y_i - \bar{y})}{\sqrt{\sum_{i=1}^{n}(x_i - \bar{x})^2}\sqrt{\sum_{i=1}^{n}(y_i - \bar{y})^2}} = \frac{\langle xy \rangle - \langle x \rangle \langle y \rangle}{\sqrt{\langle x^2 \rangle - \langle x \rangle^2}\sqrt{\langle y^2 \rangle - \langle y \rangle^2}}, \tag{S1}$$

where n is the size of the sample of joint observations of x and y, and $\langle\rangle$ denotes the sample averages of the enclosed quantities across all cells. In this case, we are considering each pair of adjacent sites, $(x, y)$, which have openness values 0 or 1. Each cell in the dataset gives another joint observation of the value of the sites. For a pair of sites, we consider the number of observations across cells of each possible openness configuration, $((0,0), (0,1), (1,0), (1,1))$, and label the proportion of such observations $p_{00}$, $p_{01}$, $p_{10}$ and $p_{11}$ respectively. Note that, since the openness values at sites $x$ and $y$ take binary values, we have:

$$\langle x^2 \rangle = \langle x \rangle = p_{11} + p_{10}, \quad \text{and} \quad \langle y^2 \rangle = \langle y \rangle = p_{11} + p_{01}. \tag{S2}$$

From Equation S1, we then have:

$$r_{xy} = \frac{p_{11} - (p_{10} + p_{11})(p_{01} + p_{11})}{\sqrt{p_{11} + p_{10} - (p_{11} + p_{10})^2}\sqrt{p_{11} + p_{01} - (p_{11} + p_{01})^2}}. \tag{S3}$$

We calculate the Pearson coefficient in this way from each site in the selected loci. We also calculate the average openness of the pair of sites, via:

$$\mu_{\text{av}} = \frac{\langle x \rangle + \langle y \rangle}{2} \tag{S4}$$

##### S1.2 Site Inhomogeneity

We consider the mean chromatin openness values at each ATAC-seq peak site within the six-site loci we selected from each dataset. Using the notation from the previous section, for a site $x$, we have a mean openness value, $\langle x \rangle$, given by:

$$\langle x \rangle = \frac{1}{n}\sum_{i=1}^{n} x_i, \tag{S5}$$

where $n$ is the number of cells in the dataset, and $x_i$ denotes the openess value (0 or 1) of the chromatin measured at site $x$ in cell $i$.

Figures S1-S3 show the mean site openness values at each site, within each locus, for all three datasets. These results indicate that, even within a single six-site locus, different ATAC-seq peak regions have significantly different average openness measurements.

#### S2 NOISE

We consider binomial dropout with probability $p_{\text{drop}}$ applied independently at each site in a locus. Figure S4 shows how this affects various distributions over three-site configurations in the Ising-like model we have described. In the case of uncorrelated (including uniformly distributed) sites, the binomial dropout effects all configurations with the same total number of open sites equally. However, for the case of correlated neighbors, ($\epsilon < 1$), the effect of dropout varies even between configurations with the same total number of open sites. For example, between $p_{\text{drop}} = 0.1$ and $p_{\text{drop}} = 0.5$, the probability of the $(0,1,0)$ configuration is reduced more than the $(0,0,1)$ and $(1,0,0)$ configurations. This is because the $(0,1,0)$ configuration ‘receives’ probability from both the $(1,1,0)$ and $(0,1,1)$ configurations, which are favored by a factor $1/\epsilon$ compared to the $(1,0,1)$ configuration.

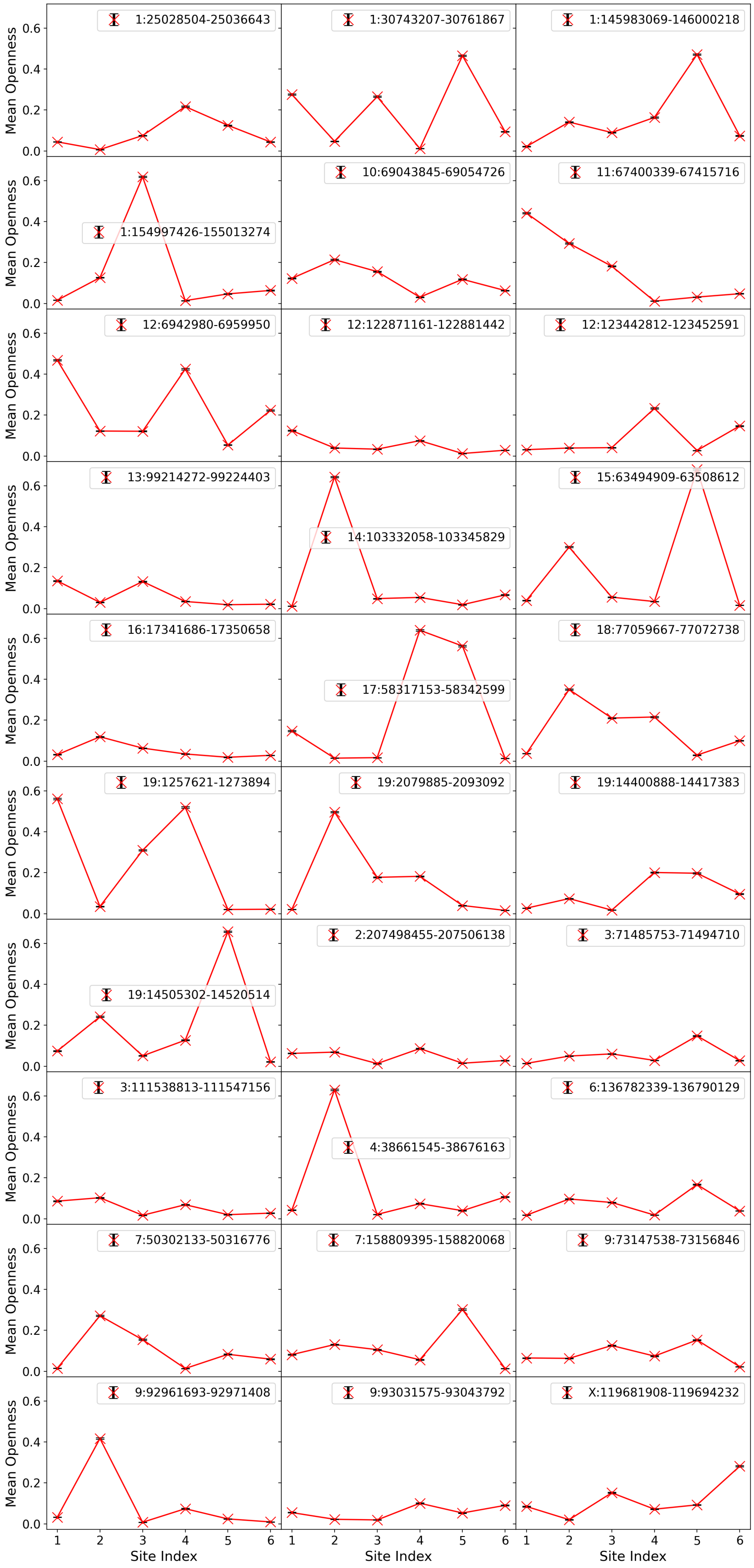


Figure S1: The mean chromatin openness for the six adjacent ATAC-seq peak sites at each of the 30 loci from the PBMC dataset.

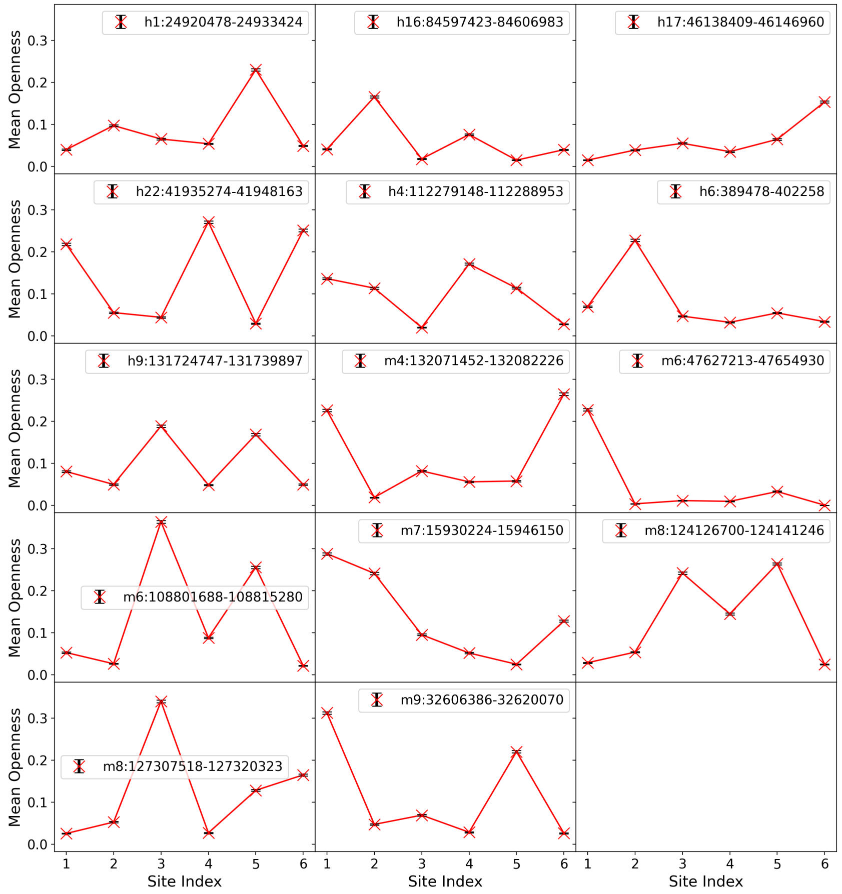


Figure S2: The mean chromatin openness for the six adjacent ATAC-seq peak sites at each of the 14 loci from the human-mouse mixture dataset.

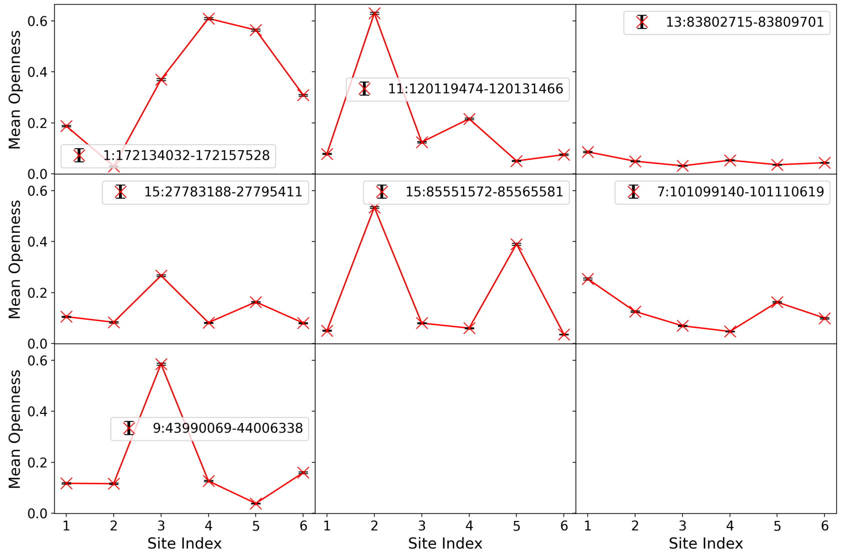


Figure S3: The mean chromatin openness for the six adjacent ATAC-seq peak sites at each of the 7 loci from the mouse cortex dataset.

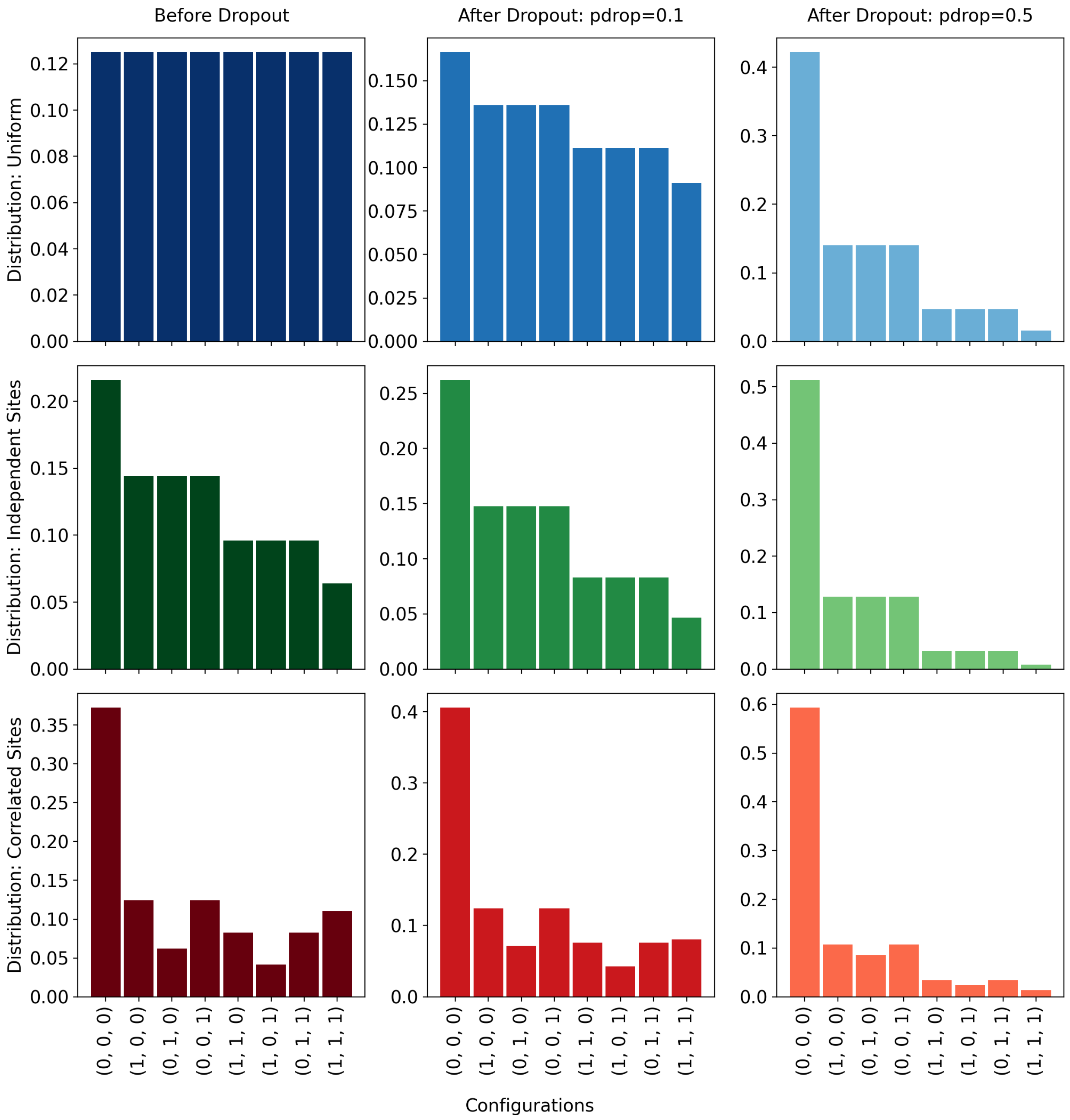


Figure S4: The effect of binomial dropout on various three-site distributions. **Top:** Uniform distribution. **Middle:** Distribution from Ising model transition matrix with independent sites. Parameters: $k_{\text{on}} = 1$ for all sites, $k_{\text{off}} = 1.5$, $\epsilon = 1$. **Bottom:** Distribution from Ising model transition matrix with correlated adjacent sites. Parameters: $k_{\text{on}} = 1$ for all sites, $k_{off} = 1.5$, $\epsilon = 0.5$

### S3 CME IMPLEMENTATION

#### S3.1 Chromatin

Working within the CME framework of (1), we consider the gene state to evolve in a Markovian manner according to the evolution equation:

$$\frac{d\boldsymbol{P}}{dt} = H\boldsymbol{P}, \tag{S6}$$

where $\boldsymbol{P}$ is a vector denoting the probability to be in each chromatin configuration. We can then encode cooperation between neighboring regions within the gene-state transition matrix, H.

As a toy example, imagine a system of only two genes, and a basis defined by the state matrix:

$$S \equiv \begin{pmatrix} 00 \\ 01 \\ 10 \\ 11 \end{pmatrix}, \tag{S7}$$

where ${S_\alpha}^i$ represents the openness of the $i^{\text{th}}$ site in the chromatin configuration indexed by $\alpha$, and takes values of 0 or 1 for closed or open chromatin respectively. In the basis we have chosen, in the first state, both genes are closed for transcription; in the second state, only the second gene is open, and so on.

We define the probability generating functions (PGF), indexed by gene state, $\alpha$:

$$G_\alpha(z,t) \equiv \sum_{\boldsymbol{m}} \boldsymbol{z}^{\boldsymbol{m}} P_\alpha(\boldsymbol{m},t), \tag{S8}$$

where $P_\alpha(\boldsymbol{m},t)$ represents the probability of the system to be in state $\alpha$ and have $m^i$ RNA counts of each transcript i, and $\boldsymbol{z}$ represents the vector of PGF arguments, $[z^1,...,z^n]$ for RNA species 1 to n. We define $\boldsymbol{z}^{\boldsymbol{m}} \equiv (z^1)^{m^1}(z^2)^{m^2}...(z^n)^{m^n}$.

Let $\boldsymbol{G}$ represent the vector of PGFs, $[G_0,...,G_{N-1}]$ for gene states 0 to N-1.

Let $\hat{B}$ be the production matrix, whose entries $\hat{B_\alpha}^i$ represent the rate of production of species $i$ in state $\alpha$. For the simplest model, where transcript i is produced at rate $b^i$ for states where gene $i$ is active and at rate zero otherwise, we have the relation: $B = S\text{diag}(\boldsymbol{b})$, for $\boldsymbol{b}$ the vector of transcription rates with components $b^i$.

Let $\boldsymbol{d}$ be a vector whose components, $d^i$, represent the decay rates of species $i$.

Following the procedure outlined in (1), we then have the full evolution equation:

$$\partial_t \boldsymbol{G} = \hat{H}^T \boldsymbol{G} + \Big[[\boldsymbol{d} \odot (1-\boldsymbol{z})]^T \boldsymbol{\partial}\Big]\boldsymbol{G} - \text{diag}[\hat{B}(1-\boldsymbol{z})]\boldsymbol{G}, \tag{S9}$$

where note that the Hadamard product between two matrices is defined via: $(A \odot B)_{\alpha\beta} \equiv A_{\alpha\beta}B_{\alpha\beta}$ for any two matrices A and B of the same dimension (here dimension (n,1) for n the number of RNA species). $(1-\boldsymbol{z})$ is the vector with components $1-z^1, 1-z^2, ..., 1-z^n$, and the second term on the right-hand side of equation S9 involves a sum over derivatives with respect to PGF variables $z^i$.

### S4 MOMENTS AND STATISTICAL PROPERTIES

The PGF evolution equation above leads to the following moments and statistical properties.

#### S4.1 Gene-State Moments

For considering moments, we introduce the vector $\boldsymbol{\pi}$, representing the steady-state probabilities of the chromatin states of the system. ($\boldsymbol{\pi}$ has the length of the entire state space, i.e. $2^n$, for n the number of sites in the locus.) In terms of the chromatin-state transition matrix, H, we have:

$$\boldsymbol{\pi} = \text{Norm}(\text{Kernel}[H^T]), \tag{S10}$$

where $\boldsymbol{\pi}$ is normalized s.t. its entries sum to one.

We then consider a state matrix S, which defines the basis we are working with, where each row of S corresponds to a possible state of the system and gives the openness of each site in the locus, where 1/0 correspond to open/closed sites respectively.

For example, a three-site system would have:

$$S = \begin{pmatrix} 0 & 0 & 0 \\ 1 & 0 & 0 \\ 0 & 1 & 0 \\ 0 & 0 & 1 \\ 1 & 1 & 0 \\ 1 & 0 & 1 \\ 0 & 1 & 1 \\ 1 & 1 & 1 \end{pmatrix}. \tag{S11}$$

In this formalism we have:

$$\langle \sigma^i \rangle = \sum_\gamma \pi_\gamma {S_\gamma}^i = (\boldsymbol{\pi}^T S)^i, \tag{S12}$$

and:

$$\langle \sigma^i \sigma^j \rangle = \sum_\gamma \pi_\gamma {S_\gamma}^i {S_\gamma}^j = [S^T \mathrm{Diag}(\boldsymbol{\pi}) S]^{ij}, \tag{S13}$$

where the $\langle\rangle$ denote expected values and $\sigma^i$ represents the openness of the $i^{th}$ region.
This gives the expression for the gene-state covariance:

$$\mathrm{Cov}(\sigma^i \sigma^j) = \sum_\beta \pi_\beta {S_\beta}^i {S_\beta}^j - \sum_{\alpha\beta} \pi_\alpha {S_\alpha}^i \pi_\beta {S_\beta}^j, \tag{S14}$$

where the variance can be obtained by setting $i = j$ in this expression.

#### S4.2 RNA Transcript Count Moments

To find moments concerning the $i^{th}$ species, we differentiate equation S9 with respect to $z^i$. This allows us to find conditional means, $\tilde{\mu}^i_\alpha$, which represent the expected number of transcripts of species $i$, given that the system is in chromatin-state $\alpha$. From the definition of $G_\alpha$ given in Equation S8, we note that:

$$\tilde{\mu}^i_\alpha = \frac{1}{\mathcal{P}_\alpha} \frac{\partial \boldsymbol{G}_\alpha}{\partial z_i}\bigg|_{\boldsymbol{z}=\boldsymbol{1}}, \tag{S15}$$

where $\mathcal{P}_\alpha$ is defined to be the probability that the system is in chromatin-state $\alpha$. We define the vector $\boldsymbol{\mathcal{P}}$ to be the vector with the $\mathcal{P}_\alpha$ as compenents. Noting that $\boldsymbol{G} = \boldsymbol{\mathcal{P}}$ for $\boldsymbol{z} = \boldsymbol{1}$, setting the LHS of equation S9 to zero and taking a derivative w.r.t. $z^i$, we reach steady-state conditional means given by:

$$\tilde{\boldsymbol{\mu}}^{\boldsymbol{i}} = [(d^i I - H^T)\hat{\Pi}]^{-1} \hat{B}^i \boldsymbol{\pi}, \tag{S16}$$

where $\hat{\Pi}$ is the diagonal matrix of $\boldsymbol{\pi}$, and $\hat{B}_i$ is the diagonal matrix of the $i^{th}$ column of B. In index notation this becomes:

$$[\hat{B}^i]_{\alpha\beta} = \delta_{\alpha\beta} S^i_\alpha b^i, \tag{S17}$$

where no summation over $\alpha$ is implied. $\tilde{\boldsymbol{\mu}}_{\boldsymbol{i}}$ is the vector whose components $\alpha$ are defined in equation S15.
For the unconditional means, $\mu^i = \sum_\alpha \tilde{\mu}^i_\alpha$, we find:

$$\mu^i = \sum_\alpha \left[(d^i I - H^T)^{-1} \hat{B}^i \boldsymbol{\pi}\right]_\alpha = \sum_\alpha \left[M^i \hat{B}^i \boldsymbol{\pi}\right]_\alpha, \tag{S18}$$

where, for ease of this and future calculations, we define:

$$M^i \equiv (d^i I - H^T)^{-1}, \text{ and } M^{ij} \equiv [(d^i + d^j) I - H^T]^{-1}. \tag{S19}$$

We also make use of the Neumann series approximation for the matrix inverse, for a matrix $\tilde{H}$ with spectral radius less than one:

$$(I - \tilde{H})^{-1} = \sum_{k=0}^{\infty} \tilde{H}^k, \tag{S20}$$

where for our purposes we define $\tilde{H}$ via:

$$H^T = d^i \tilde{H}^i, \tag{S21}$$

for a scalar, $d^i$. Then, if the spectral radius of $\tilde{H}^i/d^i < 1$, we have:

$$M^i = (d^i I - H^T)^{-1} = \frac{1}{d^i} \sum_{k=0}^{\infty} (\tilde{H}^i)^k. \tag{S22}$$

Recalling the symmetry of $H^T$ and hence $\tilde{H}$, we have:

$$\sum_{\alpha} \tilde{H}_{\alpha\beta} = 0. \tag{S23}$$

Inserting the Neumann expansion for $M^i$ in equation S18, and using the symmetry in equation S23, we see that:

$$\mu^i = \frac{1}{d^i} \sum_{\alpha} S_{\alpha}{}^i b^i \pi_{\alpha} = \frac{b^i}{d^i} \langle \sigma^i \rangle. \tag{S24}$$

For correlations between species, we use the equation:

$$\frac{1}{\mathcal{P}_{\alpha}} \frac{\partial^2 G_{\alpha}}{\partial z^i \partial z^j} = \langle m^i m^j \rangle_{\alpha}, \tag{S25}$$

noting that here and in what follows, we require $i \neq j$. Differentiating equation S9 twice, we get, in steady state:

$$0 = H^T \hat{\Pi} \tilde{\boldsymbol{\mu}}^{\boldsymbol{ij}} - d^i \hat{\Pi} \tilde{\boldsymbol{\mu}}^{\boldsymbol{ij}} - d^j \hat{\Pi} \tilde{\boldsymbol{\mu}}^{\boldsymbol{ij}} + \hat{B}^i \hat{\Pi} \tilde{\boldsymbol{\mu}}^{\boldsymbol{j}} + \hat{B}^j \hat{\Pi} \tilde{\boldsymbol{\mu}}^{\boldsymbol{i}}, \tag{S26}$$

where $(\tilde{\mu}^{ij})_{\alpha} = \langle m^i m^j \rangle_{\alpha}$ is the conditional expectation of the product of transcripts i and j given gene state $\alpha$.
Solving for the mean product vector over gene states, we get:

$$\tilde{\boldsymbol{\mu}}^{\boldsymbol{ij}} = [((d^i + d^j)I - H^T)\hat{\Pi}]^{-1} (\hat{B}^i \hat{\Pi} \tilde{\boldsymbol{\mu}}^{\boldsymbol{j}} + \hat{B}^j \hat{\Pi} \tilde{\boldsymbol{\mu}}^{\boldsymbol{i}}). \tag{S27}$$

Convolving this with the steady state probability distribution for states, we can find the unconditional expectation value of the product of two genes. Subtracting the product of the means of each gene, we arrive at the correlation between the two gene counts.

The unconditional mean product is given by:

$$\mu^{ij} = \sum_{\alpha} \left( [(d^i + d^j)I - H^T]^{-1} [\hat{B}^i \hat{\Pi} \tilde{\boldsymbol{\mu}}^{\boldsymbol{j}} + \hat{B}^j \hat{\Pi} \tilde{\boldsymbol{\mu}}^{\boldsymbol{i}}] \right)_{\alpha} = \sum_{\alpha} \left( M^{ij} [\hat{B}^i \hat{\Pi} \tilde{\boldsymbol{\mu}}^{\boldsymbol{j}} + \hat{B}^j \hat{\Pi} \tilde{\boldsymbol{\mu}}^{\boldsymbol{i}}] \right)_{\alpha}, \tag{S28}$$

with $M^{ij}$ as defined above (S19).
Converting into index notation and using the relation between B and S (S17), we find:

$$\mu^{ij} = b^i b^j \sum_{\alpha\beta} M^{ij}_{\alpha\beta} \sum_{\gamma} [S_{\beta}{}^i M^j_{\beta\gamma} S_{\gamma}{}^j + S_{\beta}{}^j M^i_{\beta\gamma} S_{\gamma}{}^i] \pi_{\gamma}. \tag{S29}$$

Using another symmetry argument related to S23 (see supplementary info S6.3.2), this is always equivalent to:

$$\mu^{ij} = \frac{b^i b^j}{d^i + d^j} \sum_{\beta\gamma} [S_{\beta}{}^i M^j_{\beta\gamma} S_{\gamma}{}^j + S_{\beta}{}^j M^i_{\beta\gamma} S_{\gamma}{}^i] \pi_{\gamma}, \tag{S30}$$

Note that, since to zeroth order in $\tilde{H}$, (i.e. in the limit of very slow switching, $k_{off}, k_{on} \ll d^i, d^j$ and $\epsilon$ not too small), we have:

$$M^{i(0)} = \frac{1}{d^i} I, \text{ and } M^{ij(0)} = \frac{1}{d^i + d^j} I, \tag{S31}$$

giving, to zeroth order in $\tilde{H}$:

$$\mu^{ij(0)} = \frac{b^i b^j}{d^i d^j}\langle\sigma^i\sigma^j\rangle, \tag{S32}$$

as expected.

From equation S28 and the expansion of $M^i$, in the case of bounded $\tilde{H}$ we have the exact expression for the covariance:

$$\mathrm{Cov}(m^i m^j) = \frac{b^i b^j}{d^i d^j}\mathrm{Cov}(\sigma^i\sigma^j) + \frac{b^i b^j}{d^i + d^j}\sum_{\beta\gamma}\sum_{k=1}^{\infty}\pi_\gamma[\frac{1}{d^j}S^i_\beta(\tilde{H}^j)^k_{\beta\gamma}S_\gamma{}^j + \frac{1}{d^i}S^j_\beta(\tilde{H}^i)^k_{\beta\gamma}S_\gamma{}^i]. \tag{S33}$$

If $\tilde{H}$ is not bounded, the series will not converge and we need to return to the expression in equation S29.

Note that, due to the structure of $\tilde{H}$, the first order term in $\tilde{H}$ always evaluates to zero (see appendix S6.3). We also show that both terms in the sum are identical, except for factors of $d^{i,j}$, giving:

$$\mathrm{Cov}(m^i m^j) = \frac{b^i b^j}{d^i d^j}\mathrm{Cov}(\sigma^i\sigma^j) + \frac{b^i b^j}{d^i + d^j}\sum_{\beta\gamma}\sum_{k=2}^{\infty}\pi_\gamma S^i_\beta(H^T)^k_{\beta\gamma}S_\gamma{}^j\left[\left(\frac{1}{d^j}\right)^{k+1} + \left(\frac{1}{d^i}\right)^{k+1}\right]. \tag{S34}$$

Again, we see that for switching which is slow compared to the RNA decay rates, i.e. $\tilde{H} << I$ for all entries of the matrix, $\mathrm{Cov}(m^i m^j) \sim \frac{b^i b^j}{d^i d^j}\mathrm{Cov}(\sigma^i\sigma^j)$, as expected.

Following the procedure above, we now consider the variance of transcript i.

$$\begin{aligned}\mathrm{Var}(m^i) &= \sum_\alpha[(2d^i I - H^T)^{-1}2\hat{B}^i\hat{\Pi}\tilde{\mu}^i]_\alpha + \mu^i - (\mu^i)^2 = \sum_{\alpha\gamma\nu}2M^{[2i]}_{\alpha\gamma}b^i S_\gamma{}^i M^i_{\gamma\nu}b^i S^i_\nu\pi_\nu + \mu^i - (\mu^i)^2 \\ &= \sum_{\gamma\nu}\frac{1}{d^i}b^i S_\gamma{}^i M^i_{\gamma\nu}b^i S^i_\nu\pi_\nu + \mu^i - (\mu^i)^2 = \frac{(b^i)^2}{d^i}\sum_{\alpha\beta}\left(S_\alpha{}^i M^i_{\alpha\beta}S_\beta{}^i\pi_\beta\right) + \mu^i - (\mu^i)^2\end{aligned} \tag{S35}$$

where we have defined matrix $M^{[2i]}$ via $M^{[2i]} \equiv (2d^i I - H^T)^{-1}$, and used the explicit decompositions of $\hat{B}$ and $\hat{\Pi}$. For the third equality we have used the argument in section S6.3.2, to give the relation $\sum_\alpha M^{[2i]}_{\alpha\gamma} = \frac{1}{2d^i}\sum_\alpha I_{\alpha\beta}$.

For the case of bounded $\tilde{H}$, this becomes:

$$\begin{aligned}\mathrm{Var}(m^i) &= \left(\frac{b^i}{d^i}\right)^2[\langle\sigma^i\sigma^i\rangle + \sum_{\gamma\beta}\sum_{k=1}^{\infty}S_\gamma{}^i\tilde{H}^k_{\gamma\beta}S_\beta{}^i\pi_\beta] + \frac{b^i}{d^i}\langle\sigma^i\rangle - \left(\frac{b^i}{d^i}\right)^2\langle\sigma^i\rangle^2 \\ &= \left(\frac{b^i}{d^i}\right)^2\left[\mathrm{Var}(\sigma^i) + \sum_{\gamma\beta}\sum_{k=1}^{\infty}S_\gamma{}^i\tilde{H}^k_{\gamma\beta}S_\beta{}^i\pi_\beta\right] + \frac{b^i}{d^i}\langle\sigma^i\rangle\end{aligned} \tag{S36}$$

#### S4.3 Correlation Propagation

We define site-site correlations for accessibility and transcript count respectively via:

$$\rho^{ij}_\sigma = \frac{\mathrm{Cov}(\sigma^i\sigma^j)}{\sqrt{\mathrm{Var}(\sigma^i)\mathrm{Var}(\sigma^j)}}, \quad \rho^{ij} = \frac{\mathrm{Cov}(m^i m^j)}{\sqrt{\mathrm{Var}(m^i)\mathrm{Var}(m^j)}}. \tag{S37}$$

We define f, the ratio between these quantities via:

$$\rho^{ij} = f\rho^{ij}_\sigma. \tag{S38}$$

Returning to matrix notation, and dropping the assumption of bounded $\tilde{H}$, we recall:

$$M^i \equiv (d^i I - H^T)^{-1}. \tag{S39}$$

$$\mathrm{Cov}(m^i m^j) = \frac{b^i b^j}{d^i + d^j}\sum_{\beta\gamma}[S_\beta{}^i M^j_{\beta\gamma}S_\gamma{}^j + S_\beta{}^j M^i_{\beta\gamma}S_\gamma{}^i]\pi_\gamma - \mu^i\mu^j, \tag{S40}$$

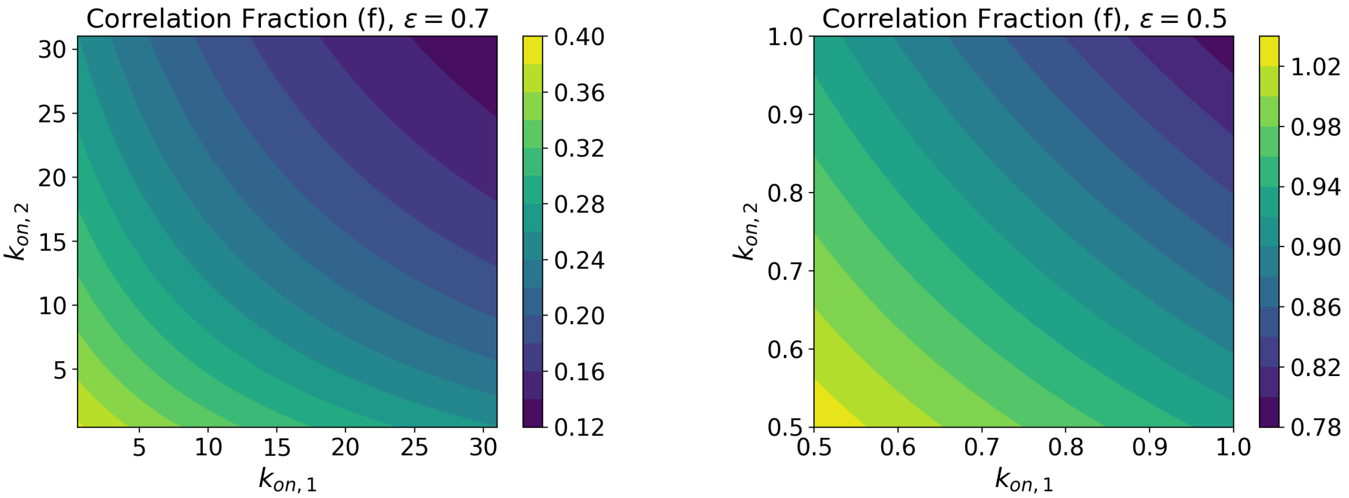


Figure S5: Ratio, f, calculated for different values of $k_{\text{on},1}, k_{\text{on},2}$ in a two-gene system. The other parameters are, **Left:** $\epsilon = 0.7, k_{\text{off}} = 30, b_1 = b_2 = 10, d_1 = d_2 = 1$, **Right:** $\epsilon = 0.5, k_{\text{off}} = 0.5, b_1 = b_2 = 10, d_1 = d_2 = 1$.

$$\text{Var}(m^i) = \frac{(b^i)^2}{d^i} \sum_{\alpha\beta} \left( S_\alpha{}^i M^i_{\alpha\beta} S_\beta{}^i \pi_\beta \right) + \mu^i - (\mu^i)^2. \tag{S41}$$

Compare these to the gene-state covariance and variances:

$$\text{Cov}(\sigma^i \sigma^j) = \sum_\alpha \pi_\alpha S_\alpha{}^i S_\alpha{}^j - \sum_{\alpha\beta} \pi_\alpha S_\alpha{}^i \pi_\beta S_\beta{}^j, \tag{S42}$$

where the variance is obtained by setting $i = j$ in this expression. An expression for f is obtained by taking the appropriate ratios of these moments:

$$f = \left[ \frac{\frac{b^i b^j}{d^i + d^j} \sum_{\beta\gamma} [S_\beta{}^i M^j_{\beta\gamma} S_\gamma{}^j + S_\beta{}^j M^i_{\beta\gamma} S_\gamma{}^i] \pi_\gamma - \mu^i \mu^j}{\sqrt{\frac{(b^i)^2}{d^i} \sum_{\alpha\beta} \left( S_\alpha{}^i M^i_{\alpha\beta} S_\beta{}^i \pi_\beta \right) + \mu^i - (\mu^i)^2} \sqrt{\frac{(b^j)^2}{d^j} \sum_{\alpha\beta} \left( S_\alpha{}^j M^j_{\alpha\beta} S_\beta{}^j \pi_\beta \right) + \mu^j - (\mu^j)^2}} \right] \left[ \frac{\sum_\alpha \pi_\alpha S_\alpha{}^i S_\alpha{}^j - \sum_{\alpha\beta} \pi_\alpha S_\alpha{}^i \pi_\beta S_\beta{}^j}{\sqrt{\sum_\alpha \pi_\alpha S_\alpha{}^i S_\alpha{}^i - \sum_{\alpha\beta} \pi_\alpha S_\alpha{}^i \pi_\beta S_\beta{}^i} \sqrt{\sum_\alpha \pi_\alpha S_\alpha{}^j S_\alpha{}^j - \sum_{\alpha\beta} \pi_\alpha S_\alpha{}^j \pi_\beta S_\beta{}^j}} \right]^{-1}. \tag{S43}$$

We show the value of f for a two-gene system with varying $k_{\text{on},1}, k_{\text{on},2}$ and two sets of remaining parameters in figure S5. The other parameters are: $b_1 = b_2 = 10, d_1 = d_2 = 1$, and $\epsilon = 0.7, k_{\text{off}} = 30$; $\epsilon = 0.5, k_{\text{off}} = 0.5$ for the left and right panels respectively.

Since RNA counts are downstream of chromatin dynamics and add an additional layer of stochasticity, we might expect that f would be constrained within the range: $0 < f < 1$. Whilst the left panel in figure S5 shows a parameter regime where this constraint is observed, the right panel provides an example where $f > 1$. The intuition behind this surprising result is explored further in section S4.4.

#### S4.4 Illustrative Toy Systems

Here we consider illustrative toy systems where the fraction, f, of transcript correlation to gene-state correlation, falls outside of the naively expected range, $0 < f < 1$. To emphasize how this can be achieved, we demonstrate the behavior of two different two-gene systems.

The first system, although un-biological, shows how Markov state transitions combined with constitutive transcript production can give transcript correlations without any correlations in the underlying chromatin openness. This corresponds to an infinite value of $|f|$.

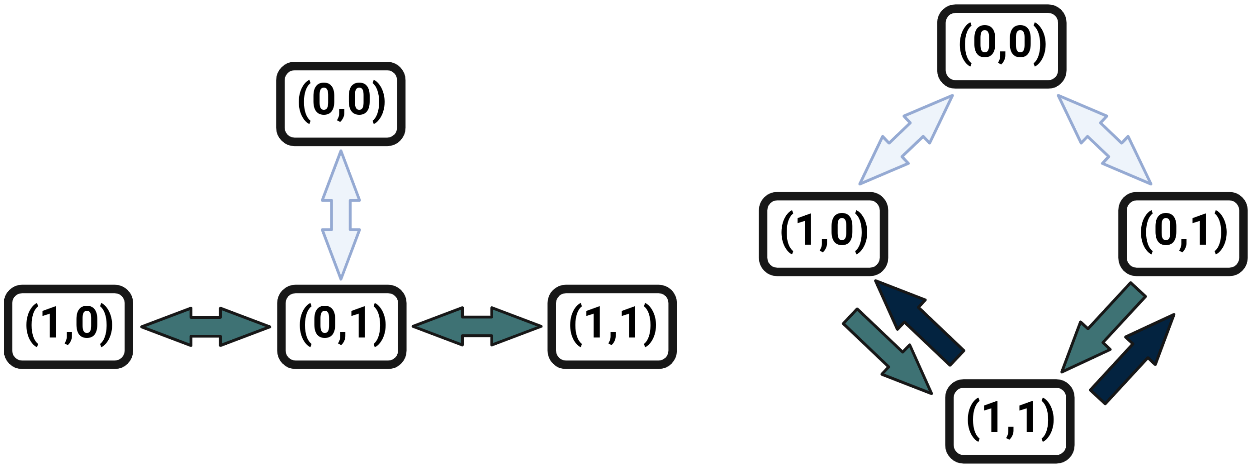


Figure S6: Simple toy systems for which the value of $f$ falls outside of the naively expected range. The states $(i, j)$ correspond to the openness of sites $i$ and $j$ in that state, where 0, 1 indicate closed and open chromatin respectively. The arrows between chromatin states represent allowed transitions, and their weights correspond to the rates of these transitions. **Left:** A system with $|f| > 1$. **Right:** A system with $f < 0$, i.e. reversed correlations between the transcript and chromatin-state levels.

The chromatin-state structure of this imagined system is shown in the left panel of Figure S6, where the chromatin-states are labeled $(i, j)$, for $i, j = 0, 1$ depending on the openness of each of the two chromatin-sites. The arrows in the diagram represent transitions between chromatin configurations, and their weight represents the rate of each transition.

We could encode such a system using the following transition matrix:

$$H = \begin{pmatrix} -\delta k & 0 & \delta k & 0 \\ 0 & -k & k & 0 \\ \delta k & k & -(2+\delta)k & k \\ 0 & 0 & k & -k \end{pmatrix} \tag{S44}$$

with $\delta < 1$, which has a uniform stationary distribution. However, due to the structure of the graph in figure S6, whilst there is no correlation between the openness of genes 0 and 1 in the steady state, there is clearly a correlation between their time averaged histories. This is because, if, for instance, gene 1 is on, that indicates that the system is in the lower half of the state graph diagram, making it more likely that it has also been in the lower half for its recent history. This means that knowing that gene 1 is on increases the probability that gene 2 has been on in the system's recent history. In our gene model, this corresponds to a correlation between transcript numbers, despite there being no correlation between the on-ness of genes 1 and 2 in steady state. The correlations resulting from simulations of this system are shown on the top row of Figure S7.

The second example system, (no longer including un-biological transitions), has reversed sign correlations (i.e. $f < 0$). We consider chromatin-state evolution defined by the transition matrix:

$$H = \begin{pmatrix} -2\delta k & \delta k & \delta k & 0 \\ \delta k & -k(1+\delta) & 0 & k \\ \delta k & 0 & -k(1+\delta) & k \\ 0 & k\chi & k\chi & -2k\chi \end{pmatrix} \tag{S45}$$

for $\delta < 1, \chi > 1$, and illustrated in the right hand panel of Figure S6. The correlations observed for such a system from simulations are shown in the bottom row of Figure S7.

#### S5 DATA PROCESSING

We processed all datasets using the snATAK pipeline (2). The pipeline provides a standardized process with a few parameters, which we list here for each dataset.

The datasets used were from 10x:

- 10k Human PBMCs, ATAC v2, Chromium Controller

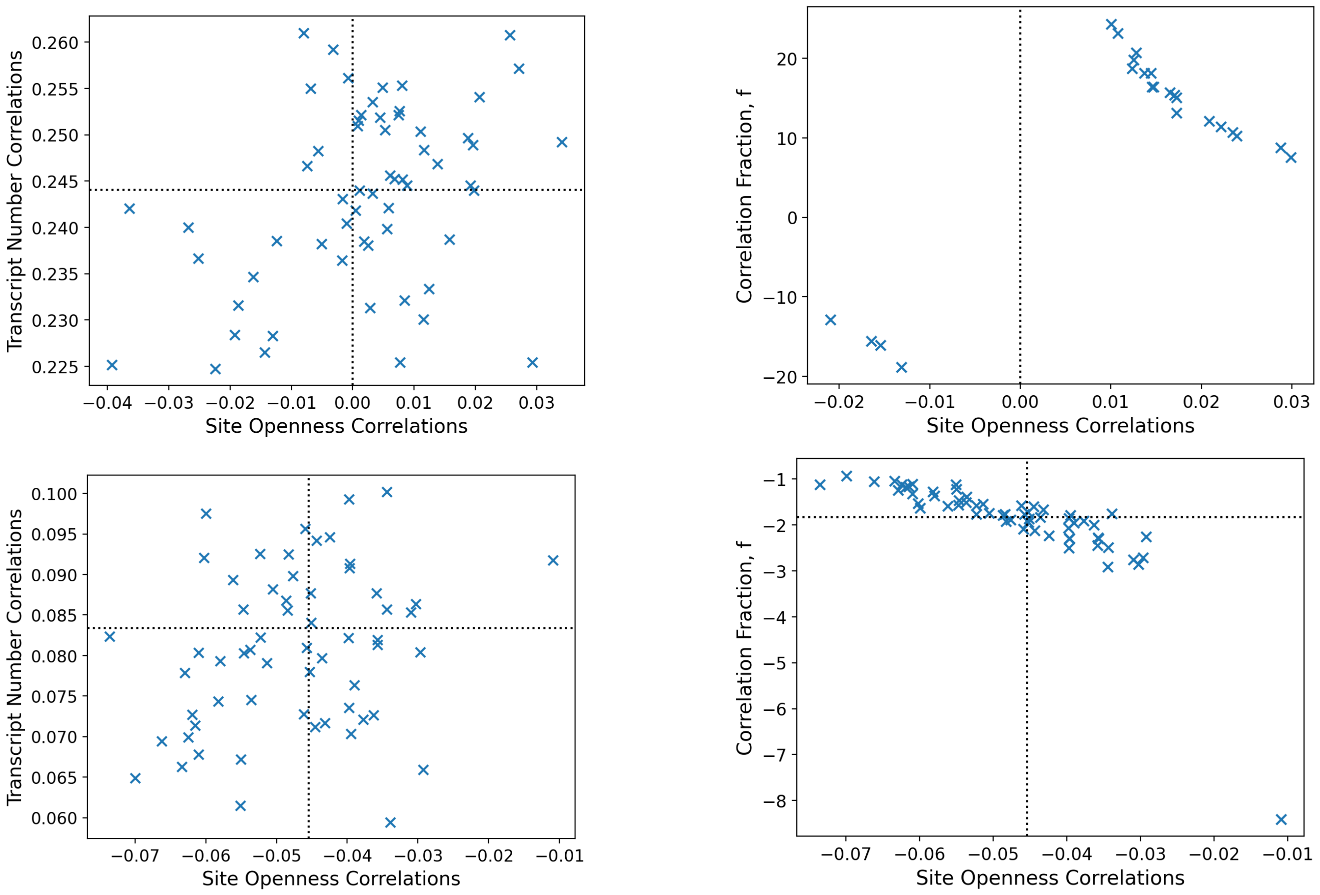


Figure S7: Correlations for simulated toy systems, with analytic solutions shown as dotted lines. **Left**: Transcript correlations versus site openness correlations. **Right:** Correlation ratio, f, versus site openness correlations. **Top:** First toy system described, with $|f| > 1$. **Bottom:** Second toy system described, with $f < 0$. The parameters used were: $\delta = 0.1, \chi = 1.2, k = 1, b_1 = b_2 = 2, d_1 = d_2 = 0.5$, and results shown are for 5000 simulated cells.

- 8k Adult Mouse Cortex Cells, ATAC v2, Chromium Controller
- 10k 1:1 Mixture of Human GM12878 and Mouse EL4 Cells, ATAC v2, Chromium Controller

To process these datasets, we used a technology string (argument option -x), which specifies the structure of the snATAK input, given by:

$$0,0,0:-1,0,0:1,0,0,2,0,0$$

indicating that there are no UMIs for this ATAC-seq data. Here files 0, 1 and 2 correspond to the R2, R1 and R3 fastq files respectively. This technology string indicates that the cell barcodes are found in the R2 file, and the paired end reads in files R1 and R3, and that the entirety of each file is used. For a whitelist we used the reverse complement of the whitelist provided by 10x. For the mixed mouse-human dataset, we first separated the human and mice cells and removed doublets.

##### S5.1 Dataset 1

#### S6 MODEL FITTING

##### S6.1 Fits at Individual Loci

Fits at the first 5 loci in dataset 1 are included in figure S8. The models are as described in the main text, and the parameters were fitting using Python's `differential_evolution` package. The algorithm was run 10 times per locus per model, to verify that the method converged closely to the same parameters each time.

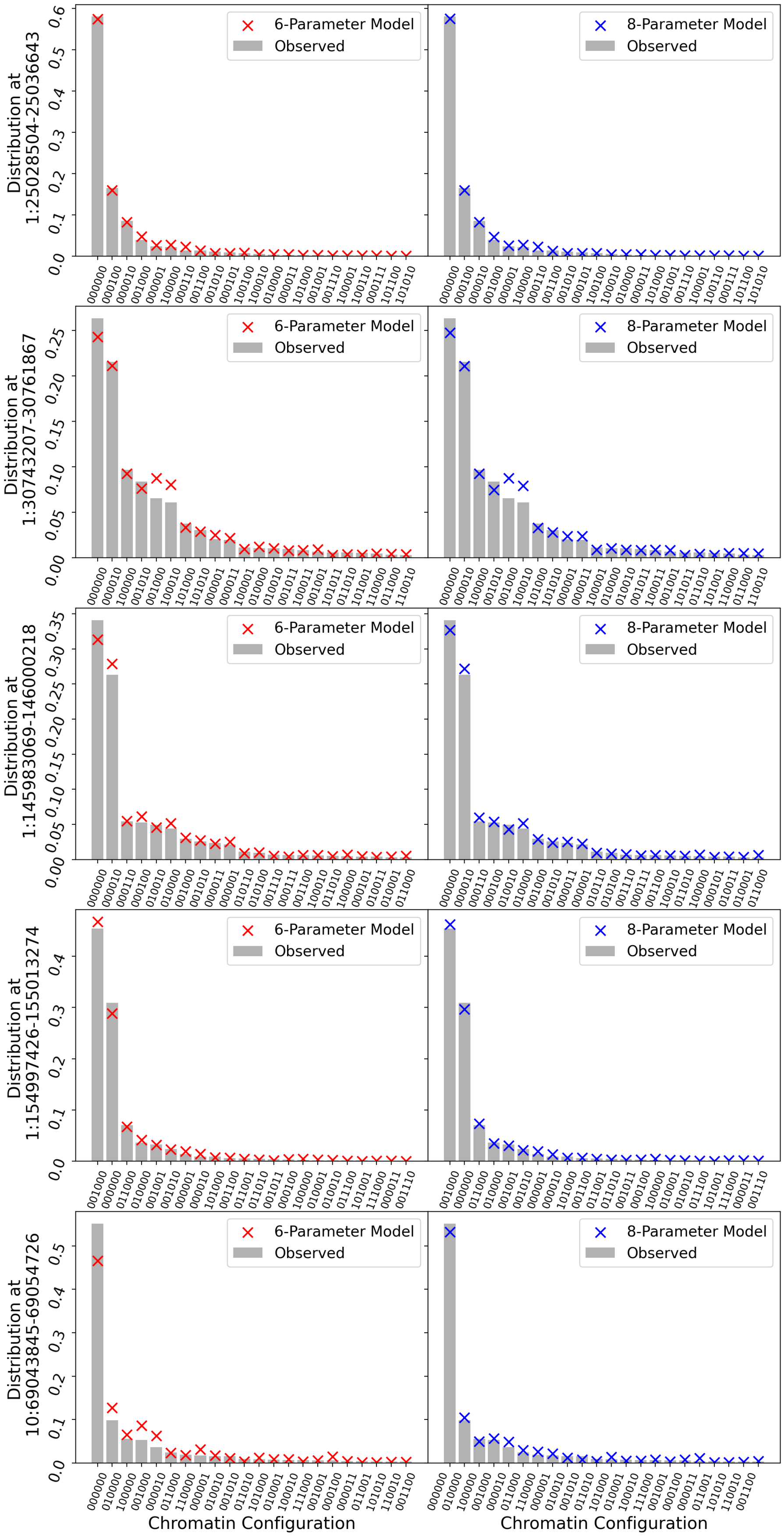


Figure S8: 6 and 8-parameter model fits to 5 loci in the human PBMC dataset. The scatter plots show the analytic distribution at the best fitting parameters, and the bar chart shows the empirical distribution.

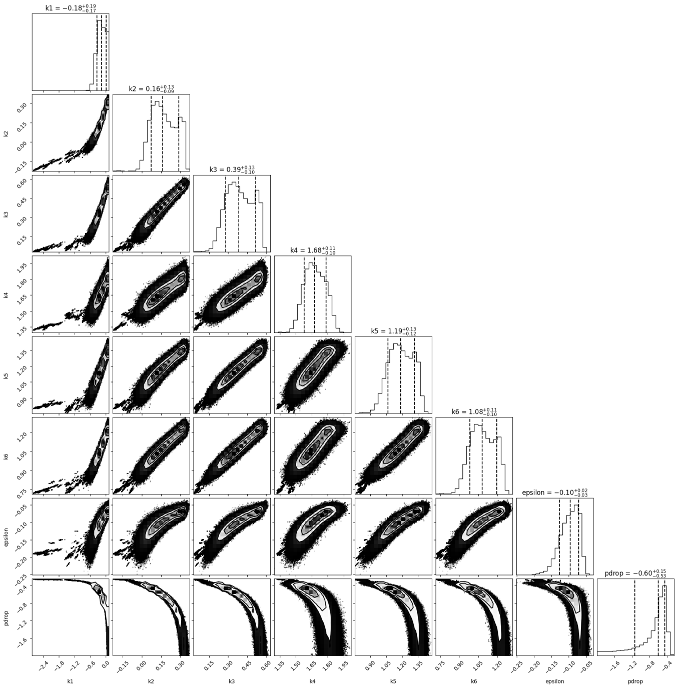


Figure S9: MCMC methods used to fit the 8-parameter model on locus 11:67400339-67415716. The most probable parameter values matched with those found using the global minimization algorithm. 512 walkers were used, with initial values of $k_{\text{on,i}} = 1$ for all sites, $\epsilon = 1$ and $p_{\text{drop}} = 0.7$. The parameters were fit in log-space.

We also fit a locus from the 10x PBMC data using MCMC (see figure S9). The parameters found by the `differential_evolution` algorithm matched the most likely values from the MCMC results.

The Bayesian Information Criteria (BIC) for the fits at each locus are shown in the main text. The BIC is defined as:

$$\text{BIC} = k\ln(n) - 2\ln\hat{L}, \tag{S46}$$

where $k$ is the number of parameters in the model, $n$ is the number of data-points, and $\hat{L}$ is the likelihood of the data under the fitted parameters.

#### S6.2 Positive Outlier: Locus 10:69043845-69054726

In the PBMC dataset, locus 10:69043845-69054726 shows a significantly larger BIC difference in favor of the 8-parameter model than any other locus. The fits of each model are shown in the bottom row of figure S8. Figure S1 (the second row) also shows that this locus seems to have mean openness values that are correlated between neighboring sites.

#### S6.3 Symmetries

##### S6.3.1 Transition Matrix Symmetry

Consider terms of the form:

$$\sum_{\alpha\beta} S_{\alpha}{}^{i} \tilde{H}_{\alpha\beta} S_{\beta}{}^{j} \pi_{\beta}, \tag{S47}$$

and recalling that $\tilde{H} \propto H^T$, we note that since the components $\tilde{H}_{\alpha\beta}$ are only non-zero for states which are connected (denoted $\alpha \sim \beta$), meaning that $\alpha$ can be reached from $\beta$ by flipping the openness of one site. Recall that $S_{\alpha}{}^{i} = 0, 1$ indicates that site i is closed/open in state $\alpha$. Since $S_{\alpha}{}^{i}$ and $S_{\beta}{}^{j}$ must be one for non-zero contributions to S47, and $\alpha \sim \beta$, we need only consider the transitions:

$$\sum_{\alpha}\sum_{\beta\sim\alpha} S_{\alpha}{}^{i} S_{\beta}{}^{j}\left(S_{\beta}{}^{i} S_{\alpha}{}^{j} + (1 - S_{\beta}{}^{i})S_{\alpha}{}^{j} + (1 - S_{\alpha}{}^{j})S_{\beta}{}^{i}\right)\tilde{H}_{\alpha\beta}\pi_{\beta}, \tag{S48}$$

where we have divided the sum into transitions between connected states, i.e. excluding those where both sites i and j flip between states $\alpha$ and $\beta$. The first term represents transitions between states where both sites i and j are open in both $\alpha$ and $\beta$, the second from both open to i open and j closed, and the last from i closed and j open to both open. Then, noting the Markovian properties of $H^T$ and $\pi$, we have that:

$$\sum_{\alpha,\alpha\sim\beta,\alpha\neq\beta} \tilde{H}_{\alpha\beta}\pi_{\beta} = \sum_{\alpha,\alpha\sim\beta,\alpha\neq\beta} \tilde{H}_{\beta\alpha}\pi_{\alpha}, \tag{S49}$$

allowing us to re-write the final term of equation S48 as:

$$\sum_{\alpha}\sum_{\beta\sim\alpha} S_{\alpha}{}^{i} S_{\beta}{}^{j}(1 - S_{\alpha}{}^{j})S_{\beta}{}^{i}\tilde{H}_{\alpha\beta}\pi_{\beta} = \sum_{\alpha}\sum_{\beta\sim\alpha} S_{\alpha}{}^{i} S_{\beta}{}^{j}(1 - S_{\alpha}{}^{j})S_{\beta}{}^{i}\tilde{H}_{\beta\alpha}\pi_{\alpha} = \sum_{\alpha}\sum_{\beta\sim\alpha} S_{\beta}{}^{i} S_{\alpha}{}^{j}(1 - S_{\beta}{}^{j})S_{\alpha}{}^{i}\tilde{H}_{\alpha\beta}\pi_{\beta}, \tag{S50}$$

giving the total expression:

$$\sum_{\alpha} S_{\alpha}{}^{i} S_{\alpha}{}^{j} \sum_{\beta\sim\alpha}\left(S_{\beta}{}^{j} S_{\beta}{}^{i} + (1 - S_{\beta}{}^{i})S_{\beta}{}^{j} + (1 - S_{\beta}{}^{j})S_{\beta}{}^{i}\right)\tilde{H}_{\alpha\beta}\pi_{\beta} = \sum_{\alpha} S_{\alpha}{}^{i} S_{\alpha}{}^{j} \sum_{\beta\sim\alpha} \tilde{H}_{\alpha\beta}\pi_{\beta} = 0, \tag{S51}$$

where the second equality comes from the fact that $S_{\alpha}{}^{i} S_{\alpha}{}^{j}$ selects for transitions into states with both sites i and j open. The connected origin states $\beta$ must either have both states i and j open, or exactly one of the i and j sites open. Hence the sum in terms of $S_{\beta}{}^{i,j}$ expresses all of the non-zero transitions between states $\beta$ and $\alpha$, making the expression redundant with the specification that all $\beta \sim \alpha$. The third equality comes from the symmetry of $\tilde{H}$ (S23), and the fact that only $\tilde{H}_{\alpha\beta}$ are only non-zero for $\alpha \sim \beta$.

##### S6.3.2 Inverse Matrix Symmetry

Using the transition matrix symmetry:

$$\sum_{\alpha} \tilde{H}_{\alpha\beta} = 0, \tag{S52}$$

we consider a matrix of the form:

$$M \equiv (I - \tilde{H})^{-1}. \tag{S53}$$

Note that, by the definition of M:

$$(I - \tilde{H})M = I, \tag{S54}$$

and that, also:

$$(I - \tilde{H})M = M - \tilde{H}M. \tag{S55}$$

Considering the RHS of equation S55 component-wise, and summing over $\alpha$, we note that:

$$\sum_{\alpha}[M_{\alpha\beta} - \sum_{\gamma} \tilde{H}_{\alpha\gamma} m_{\gamma\beta}] = \sum_{\alpha} M_{\alpha\beta} - \sum_{\gamma}\sum_{\alpha} \tilde{H}_{\alpha\gamma} m_{\gamma\beta} = \sum_{\alpha} M_{\alpha\beta}, \tag{S56}$$

where for the last equality we have used the symmetry of $\tilde{H}$ (S52). Then, equating this with the component-wise version of equation S54, we arrive at:

$$\sum_{\alpha} M_{\alpha\beta} = \sum_{\alpha} I_{\alpha\beta}, \tag{S57}$$

as desired. □
